# Helix recesses boost coral larvae settlement and survival

**DOI:** 10.1101/2024.11.14.623685

**Authors:** Jessica Reichert, Hendrikje Jorissen, Crawford Drury, Joshua R. Hancock, Corryn Haynes, Allison Nims, Lomani H. Rova, Nina M. D. Schiettekatte, the R3D Consortium, Joshua S. Madin

**Affiliations:** Hawaiʻi Institute of Marine Biology, University of Hawaiʻi at Mānoa, Hawaiʻi, Kāneʻohe, USA; University of Hawaiʻi at Mānoa, Hawaiʻi, Kāneʻohe, USA

**Keywords:** 3D printing, artificial reefs, coral settlement habitat, hydrodynamics, Kāneʻohe Bay Hawaiʻi, scalable restoration

## Abstract

The worldwide decline of coral reefs, driven by climate change and local stressors, demands new, scalable restoration approaches. Coral larvae offer significant potential for reef recovery, as a single coral colony can release millions of offspring, with almost all larvae dying before finding a suitable habitat. Building upon an iterative design process, incorporating a gradient of sizes and angles based on larval settlement preferences observed in nature, we developed and tested seven 3D-printed settlement module designs, proposed to enhance coral larvae habitat. We studied the settlement and survival of coral larvae on the settlement modules integrating these designs and adjacent reef structures in Kāneʻohe Bay over one year. Helix recesses dramatically outperformed other structural features, increasing settlement by ∼80-fold and post-settlement survival over a year by 20-50-fold compared to control modules. In contrast to natural reef substrates, settlement on modules with helix recesses increased by ∼70-fold. We identified the recess dimensions and light levels preferred as settlement habitat. In a parallel tank experiment, we explored the impacts of hydrodynamics on the settlement and survival of *Montipora capitata* larvae on modules with helix recesses. We found that settlement was more pronounced under high-flow conditions, suggesting a crucial role of micro-scale hydrodynamics in entraining settling corals. These findings highlight the potential of helix recesses to significantly improve early coral recruitment, a critical bottleneck in reef restoration. By integrating these structures into artificial reefs and coastal infrastructure, our approach offers an innovative, scalable, and cost-effective solution to enhance reef resilience and accelerate ecosystem recovery in a changing ocean.

## 1 Introduction

Coral reefs worldwide are experiencing an unprecedented decline, primarily driven by climate change and local anthropogenic stressors, such as increasing temperatures, acidification, and pollution. Global reef cover has already declined by 30–50% in recent decades, with some regions, such as the Caribbean, having lost over 80% of their coral cover, and a third of reef-building coral species are considered at risk of extinction (Carpenter et al., 2008; Hughes et al., 2017). According to the Intergovernmental Panel on Climate Change, even under a 1.5°C warming scenario, 70–90% of coral reefs are projected to decline (IPCC, 2023). These combined impacts threaten critical ecosystem services, including shoreline protection, food security, and tourism, and have left many reefs with limited potential for natural recovery (Woodhead et al., 2019). Lowering carbon emissions is our best chance to mitigate coral reef decline, however, local efforts can support degrading ecosystems through restoration approaches (Randall et al., 2020; Suggett and van Oppen, 2022). Despite some successful efforts to restore reefs, the overall scale of reef restoration has not kept pace with the widespread and rapid decline of coral reefs. This is because the effectiveness, scalability, and costs of restoration methods remain challenging (Boström-Einarsson et al., 2020; Hein et al., 2017). Restoration approaches often focus on transplanting reared or fragmented coral colonies to degraded reefs (Boström-Einarsson et al., 2020). These methods have been successful on local scales (< 1 km^2^), particularly in the Caribbean and Indo-Pacific, but their propagation is labor-intensive, involves sacrificing healthy coral colonies as donor stock, and the associated high costs remain a significant barrier to large-scale restoration success. A more cost-effective and feasible propagation method is to use sexual propagules obtained during spawning (Randall et al., 2020). A few coral colonies can release millions of individuals during a single spawning event (Madin et al., 2016). Therefore, restoration approaches increasingly collect and relocate coral larvae to degraded reef areas, where natural recovery is too slow or limited (Cruz and Harrison, 2017; Harrison et al., 2021; Suzuki et al., 2020). This strategy aims to promote larvae settlement and support population recovery on damaged reefs by enhancing the supply of competent larvae. Other approaches focus on collecting and settling larvae onto seeding units that are then outplanted to target areas (Chamberland et al., 2017). Because larvae are produced sexually, this method also supports high genetic diversity comparable to that found in natural populations, making it a promising strategy for enhancing long-term reef resilience (Zayasu and Suzuki, 2019).

The process of shifting from motile planula larvae to sessile polyps includes the selection of a suitable substrate, the attachment to it, and metamorphosis (Randall et al., 2020). While the primary mode of transportation for planktonic larvae is through ocean currents, the larvae can modulate their vertical position in the water column to get in contact with the reef (Tay et al., 2011). Once they are in proximity to the reef, coral larvae can detect a variety of physical, chemical, and biological cues, which guide them to specific locations and influence settlement (Edmunds, 2023). Biochemical cues from certain crustose coralline algae (CCA) or bacteria are considered strong settlement triggers (Gómez-Lemos et al., 2018; Harrington et al., 2004; Jorissen et al., 2021). Furthermore, material composition, surface topography, and water flow can affect larvae entrapment and settlement (Gysbers et al., 2024; Hata et al., 2017; Levenstein et al., 2022b, 2022a; Whalan et al., 2015). Overall, coral larval settlement preferences are governed by a trade-off between the need for light to fuel growth and the risk of exposure. Light enables rapid early growth, allowing corals to overcome size-escape thresholds, typically reached once recruits grow to at least 3–5 mm in diameter or are a few months old, at which point mortality from predation, grazing, and competition declines significantly (Doropoulos et al., 2016). However, high-light environments also increase the risk of overgrowth by competitively superior algae. This trade-off is likely to vary among species depending on their life history traits, such as growth rate, reproductive mode, or tolerance to environmental stressors. One successful strategy to balance this trade-off is for larvae to settle in crevices and near underside edges, protecting them from predation, grazing, and strong physical impacts while still receiving sufficient light to grow (Cruz and Harrison, 2017). However, after settlement, coral recruits undergo an early-life bottleneck, and only a fraction of the larvae that initially settled survive (Martinez and Abelson, 2013; Nozawa and Harrison, 2008). Thus, improving early life survival can have a disproportionately large impact on population dynamics and ultimately coral cover, presenting a scalable tool for coral restoration efforts.

A novel approach to boost restoration efforts is the integration of settlement habitat for coral larvae into coastal structures, such as engineered breakwaters and artificial reefs (AR); a similar vein to creating habitat for other reef organisms to restore biodiversity (Harrison et al., 2021; Lima et al., 2019; Loke et al., 2015). Different methods have been applied to build artificial reefs, deploying, for example, modular concrete structures like *MARS* or hexagonal rod constructions covered with resin and coral sand like *Reef Stars* (Higgins et al., 2022). Recently, 3D printing has been applied as a versatile method for creating modules to rebuild the structural complexity of reefs (Berman et al., 2023; Levy et al., 2022). Ceramic 3D printing allows the creation of varied surface textures, crevice depths, and spatial arrangements, enabling exploration of how microhabitat features influence coral larval settlement and survival. It also facilitates rapid prototyping and testing of diverse designs to identify optimal microhabitat characteristics. However, it remains unclear which structural elements and microhabitat features should be integrated into artificial reefs to maximize settlement and survival of coral recruits. Once identified, these key features can then be incorporated into larger-scale marine engineering projects, such as breakwaters and seawalls, to support scalable restoration approaches.

Inspired by observations of concentrated larval settlement in recesses and edges on natural reef, recent work discovered a distinct preference of coral larvae for helix (i.e., angular) recesses when compared to other recess types (Schiettekatte et al. in prep). These helix structures consist of spiral recesses integrated into conical modules, creating a gradient of angled microhabitats that vary in depth, width, and exposure, designed to mimic the structural complexity of natural reef surfaces. The goal of our research was to develop a spectrum of helix designs that capture a gradient of recess sizes, angles, and helix weaves to narrow down the properties of settlement substrate preferred by coral larvae. Specifically, we I) design and compare different structural features that can be integrated into modules as settlement structures for coral larvae, II) identify the properties and dimensions of the most successful structural features, and III) assess the influence of hydrodynamics on the settlement and survival of coral recruits.

## 2 Materials and Methods

### 2.1 Experimental design

To identify structural features promoting coral settlement and survival to be applied in reef restoration or engineering structures, we developed a set of adaptable modules in an iterative design process and tested their performance in a field experiment. Larvae settlement modules were designed as conical shapes incorporating a gradient of sizes and angles in seven designs (Figure 1A). In the field experiment, these larvae settlement modules were deployed at two locations in the field (reef 13 and Moku o Loʻe) in proximity to patch reefs in Kāneʻohe Bay, Oʻahu, Hawaiʻi (Fig. 1E−G, n ≥ 5 per shape and location, see Table S1 for detailed number of replicates). Recruitment on settlement structures was compared to recruitment on natural reef substrate in the Barrier Reef in Kāneʻohe Bay.

**Figure 1:**
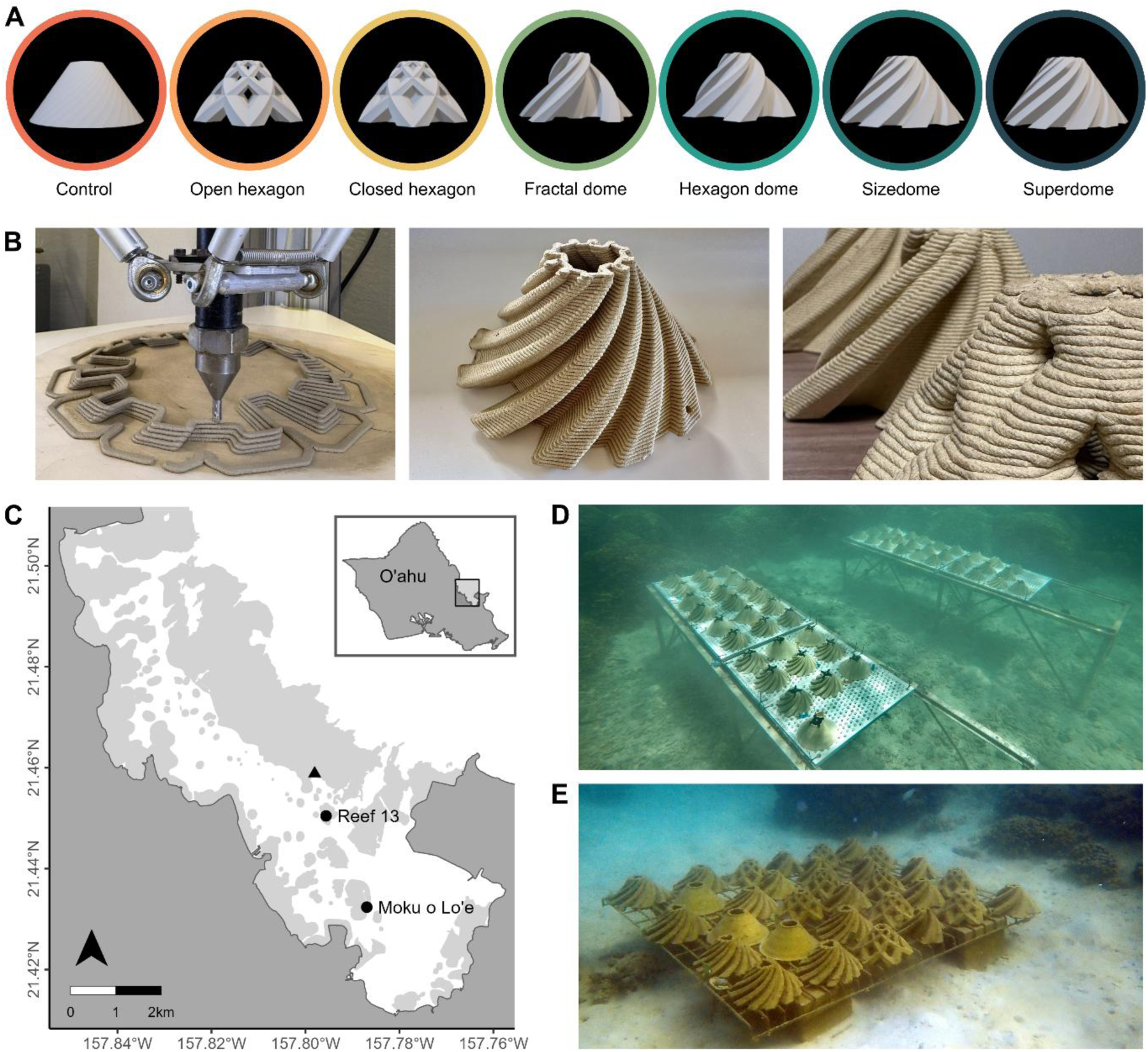
Larvae settlement module designs and production. (A) Seven different designs (control, open hexagon, closed hexagon, fractal dome, hexagon dome, sizedome, and superdome) with different structural features were compared. (B) The conical larvae settlement modules were built with a 3D clay printer. The close-up shows the horizontally caved microscale structure on the surface of the fired modules, created through the 3D printing process. (C) Map of Kāneʻohe Bay, Oʻahu, Hawaiʻi, showing the study location patch reef 13 and Moku o Loʻe. The triangle indicates the location in the barrier reef, where natural recruitment was assessed for comparison. Light grey areas show the presence of reefs. The settlement modules were deployed at underwater tables at (D) reef 13 and (E) Moku o Loʻe.

To study the impact of different flow speeds on coral settlement and survival, we performed a tank experiment. We first assessed the effects of flow on larvae settlement. For this, larvae settlement modules of superdome and control shapes were seeded with coral larvae under three different flow speeds: no, low, and high (0, 10, and 15 cm s-1, respectively). Second, we assessed the impacts of flow on recruit survival. For this, the seeded settlement modules were exposed to two different flow speeds: low flow and high flow (10 and 15 cm s-1, respectively). The seeded settlement modules were distributed equally in two tanks (n = 12 superdomes and n = 9 control domes), originating from the three seeding treatments.

Larvae settlement and recruit survival were monitored in both field and tank experiments, one week after spawning of *M. capitata* (06.07.2023), and early-life survival two and six months (30.08.2023 and 15.01.2024, respectively) after seeding. In the field, an additional timepoint was added at approximately one year after settlement (23.05.2024), before the next anticipated spawning of *M. capitata*.

All activities involving coral reef organisms, including the deployment of settlement structures and collection of coral larvae, were conducted under the Hawaiʻi Institute of Marine Biology Special Activity Permit 2024-45 and in full compliance with institutional, national, and environmental research regulations.

#### 2.1.1 Design and production of coral larvae settlement modules

Modules all shared the same base dimensions and had a diameter of 20 cm and a height of 10 cm. Two designs included holes (‘open hexagon’) or pockets (‘closed hexagon’) of different sizes (i.e., 10x12 mm, 16x24 mm, and 26x36 mm diameter). Four designs included various kinds of helix recesses (‘hexagon dome’, ‘fractal dome’, ‘sizedome’, and ‘superdome’), which decreased in size towards the top and inclined at a constant angle of 45° around the module. The ‘hexagon’ shape held six open recesses of 24−5 mm depth and 48−9 mm width (bottom to top) framed by wide-angled sidewalls (120°). The ‘fractal dome’ shape was designed to provide three exposed, sheltered, and cryptic recesses, each, of 17−3 mm depth and 32−5 mm width (bottom to top), with the same wide-angled walls (120°) as the hexagon control. The ‘sizedome’ held ten narrow recesses of 20−5 mm depth and 31-8 mm width (bottom to top), framed by narrow-angled sidewalls (100°). The ‘superdome’ held 12 narrow recesses of four different depths (10−2 mm, 20−5 mm, 30−8 mm, and 40−11 mm, referring to as depths 1, 2, 3, and 4 respectively), all with 30−6 mm width (bottom to top), framed by narrow-angled sidewalls (98°). All modules were designed as hollow cones with an open hole to ease production, handling, and attachment. As a control shape, a flat cone sharing the same base dimensions was created.

All shapes were designed in Blender v3.4. Their planar circular base design was replicated in 200 lines, stacked on top of each other, scaled to 30% of the original size, and rotated by 60 degrees (from bottom to top) so that a conical shape with equally inclining angles of the recesses was created by bridging the edge loops of the lines. For two designs, cones were mirrored and combined to create holes (from six basic hexagon shapes) and pockets (from the hexagon design, a circular shape with recesses). The resulting meshes were exported as .stl files and converted to a 3D print path using the slicing software Simplify3D. Modules were printed with the Delta Wasp 40100 3D printer (Wasp, Italy) in clay (mid-fire pottery clay, Soldate 60, Aardvark Clay and Supplies, United States), a commercially available mid-fire ceramic formulated for large-scale applications and 3D printing. The clay feed material was prepared using a pugmill (Pugmill/Mixer NVS-07, Nidec, Japan). Modules were printed in 110 % of the target size to account for the shrinking of the material, as hollow conical shapes (setting: spiral vase mode). The material was extruded with a 4 mm diameter nozzle, which created a microstructure of horizontal layers on all modules (Figure 1B). Printed modules were air-dried in a stable, air-conditioned environment for 5−10 days until they were bone-dry and then fired in a kiln (Jupiter Sectional Kiln, L&L Kilns, United States) at ∼2030 °F (≈1100 °C), held for 2 h (setting: cone 8). No surface treatment or glazing was applied after firing and modules were used in their natural, unglazed form, resulting in an open porous ceramic surface with fine horizontal ridges created by the spiral toolpath of the 3D printing process (Figure 1B).

#### 2.1.2 Deployment of field experiment

Reef 13 is a rather exposed patch reef with medium flow, located at the channel of Kāneʻohe Bay, a known pathway for larval dispersal. The site at Moku o Loʻe is a more sheltered, shallow water environment, with higher daily temperature fluctuations and lower water flow. The contrasting environmental conditions between the two sites allowed us to assess the performance of settlement structures across a natural gradient of reef habitats. The modules were attached to tables one week before coral spawning to reduce the potential for initial leaching of trace metals reported in newly fired unglazed ceramics and allow for a biofilm to form on the surface. A temperature logger (HOBO Pendant MX2201 Water Temperature Data Logger, HOBO, USA) was attached to each table, logging data at 5-minute intervals (Figure S1).

#### 2.1.3 Tank setup of flow experiment

The experiment was conducted in three 230 L tanks (180 cm x 56 cm x 23 cm), with a different flow speed treatment established in each tank. The tanks were equipped with two wavemaker pumps each (Nero 7, Aquaillumination, USA), positioned at opposite short sites. The flow was established as an oscillating wave (alternating from full intensity to no intensity) with an interval of 10 seconds, estimated at 0, 10, and 15 cm s^-1^, respectively. The flow speeds were selected to reflect low to moderate flow conditions commonly found on coral reefs (Roik et al., 2016). These levels were deliberately kept within a conservative range, to avoid potential damage to the coral larvae due to mechanical risks from the propeller pump. Larvae settlement modules of superdome and control shapes (n = 8 and n = 6 per treatment, respectively) were placed in tanks at equal flow (10 cm s^-1^) one week before the seeding to reduce the potential for leaching chemicals and allow for biofilm formation on the surface. Then, flow speeds were adjusted to the treatment conditions, and larvae were seeded (for details on collection and seeding, see sections 2.3.2 and 2.3.3).

#### 2.1.4 Collection and rearing of coral larvae and seeding of settlement modules in tanks

Coral larvae were collected during the mass spawning of *M. capitata* on June 18, 2023, from near Reef 11 in Kāneʻohe Bay, Oʻahu, Hawaiʻi following established best practices (Hancock et al., 2021). Larvae were returned to HIMB, fertilized and reared over 4 days to settlement competency at ambient temperatures in 1 µm filtered seawater following established best practices (Rahnke et al., 2022).

Coral larvae were added to the tanks 3 days after spawning. Specifically, 3 L larvae in suspension (concentration: 10.2 ± 1.4 larvae per mL, n = 3 measurements) were scooped out of a stirred, homogenized larvae-rearing bucket, and added to each tank. Larvae were added in the afternoon (5:30 pm) and left in the tanks for 24 h to allow for settlement on the modules. Water inflow was turned off during the first 18 h to facilitate an undisturbed water flow within the tanks. For the following 6 h, the tanks were equipped with 65 µm mesh sieves at the outflow to reestablish water inflow during the daytime but retain larvae within the tank. Sieves were gently rinsed every hour using a pipette to bring adhering larvae back in suspension. To ensure that the pumps of the flow treatments had no negative impact on the survival of the coral larvae, the number of dead larvae was assessed 18 h after seeding. We found no significant difference between the three tanks (Pairwise Wilcoxon test, p > 0.005).

#### 2.1.5 Water parameters and tank maintenance

The tank experiment was conducted at the Hawaiʻi Institute of Marine Biology (HIMB). The tanks were connected to a permanent saltwater supply from the reef and were provided with filtered saltwater (Pentair Triton II, TR60 sand filter, Pentair Aquatic Eco-Systems Inc., USA). Water parameters of the tank system were measured every other week using a multiparameter photometer (H197115 Hanna Labs Marine Master Multiparameter Photometer, Hanna Instruments, USA) in alternating tanks (salinity: 33.9 ± 0.4, magnesium: 1381 ± 105 ppm, phosphate: 0.2 ± 0.1 ppm, nitrate: 0.33 ± 0.35 ppm, and nitrite: 1 ± 3 ppb). Additional water parameters were measured at the beginning and the end of the experiment (calcium: 476 ± 25 ppm, ammonia: 0.03 ± 0.02 ppm, alkalinity: 6.4 ± 0.1 dKH, and pH: 7.8 ± 0). Temperature was logged with HOBO data loggers positioned in each tank, logging in intervals of 5 min (Figure S2). To avoid tank effects, modules and flow conditions were switched between tanks every other week.

### 2.2 Assessment of total recruit numbers and densities

The total numbers of recruits were assessed on all settlement modules. For size- and superdomes, the numbers per recess and recess part were additionally assessed (outside, exposed edge, inside, cryptic edge, details see results Figure 3A). Coral recruits were detected through fluorescence, using underwater blue lights (Sola Nightsea Blue Light, Light & Motion, USA) and blue-light-blocking glasses (Amazon Basics, Amazon USA). Total numbers were converted to recruit densities by standardizing to the surface area of the module. For size- and superdomes, numbers in recess parts were standardized to the respective area of the recess parts. To determine the surface areas of the modules, one dome per design was 3D scanned using the HandySCAN 3D with VX elements (Creaform, Canada) at a resolution of 0.5 mm.

Recruitment on settlement structures was compared to recruitment on natural reef substrate in the Barrier Reef in Kāneʻohe Bay after two months. There, coral recruitment was assessed in 32 0.5x0.5 m quadrants. A grid of 64 quadrats (0.5 × 0.5 m each) was placed on a representative, flat section of the barrier reef crest to ensure substrate consistency and image clarity for photogrammetry. From these, 32 quadrats were randomly selected for coral recruit counts. Coral recruits were surveyed using underwater fluorescence detection, and substrate composition and 3D structure were quantified via photogrammetry. Coral recruits were counted using underwater blue lights and blue-light-blocking glasses. Percent cover of surface compositions (CCA, macroalgae, live coral, and coral rubble/sand) was estimated visually. The 3D structure of the surveyed area was documented using photogrammetry following established procedures (Pizarro et al., 2017). Briefly, the reef was photographed following a spiral pattern from a center point, collecting ∼2500 photos, together with depth and GPS coordinates. Orthomosaics were created in Agisoft Metashape photogrammetric software (Roach et al., 2021). Coral recruitment was standardized to the area of available settlement substrate. For this, the area of live coral, macroalgae, and sand was subtracted from the quadrat areas, and the remaining area of the quadrants was translated into the surface area via rugosity (Schiettekatte et al., 2025).

### 2.3 Documentation and modeling of recruit position and recess properties

Recruit position was assessed in a subset of recruits on settlement modules of the superdome shape in the field experiment at reef 13. The recess width at locations with recruits were documented in three superdomes: four recesses each at one week, twelve recesses each at two months, six months, and one year – compensating for the initial high post-settlement mortality. The recess depth and light level at which the recruit was found were modeled as follows. Recess widths and depths were assessed at five positions for each recess depth on two printed and fired superdomes: the base of the module, 2 cm height, 4 cm height, 6 cm height, and the top of the module. The relationships between width and depth were determined for each recess (linear model, Figure S3A). The superdome modules provided a recess width-depth space of 0.9–1.3 cm width and 0.7–1.3 cm depth (Figure S3B). Light levels were measured (LI-250A Light meter, LI-COR, USA) at four positions inside the recesses, excluding the top of the module: the base of the module, 2 cm height, 4 cm height, and 6 cm height. For this, the modules were positioned under an artificial light source with ∼200 µmol m^-2^ s^-1^ photosynthetically active radiation (PAR) at the top of the modules to allow for an equal comparison under moderate light exposure, comparable to levels found at the reef. The relationship between light levels and recess width was determined for each recess depth (1, 2, 3, and 4 cm) using general additive models (GAMs) with spline-based smoothed regressions in the gam function of the R package mgcv (Wood, 2017). These were then used to predict light levels at the positions where the recruits were found. Additionally, light levels of other shapes were measured at potential settlement areas, i.e., inside the recesses at the same heights (hexagon dome, fractal dome, and sizedome) or inside the holes (open and closed hexagon), respectively. For the control dome, light levels were assessed at the same heights at three positions around the module.

### 2.4 Measurement of coral growth, biomass, and genus

Coral growth and genus were assessed in larvae settlement modules of control and superdome shape one year after spawning at both locations the field experiment in the same subset assessed for recruit position. For this, coral length and width were measured, and surface area was calculated assuming an oval shape. Coral length, width, and recruit height were used to calculate the coral volume, approximated as a cylinder. The coral genus was assigned visually after one year, where possible.

### 2.5 Statistical analysis

We performed all data processing and analyses in the R statistical environment (v.4.3.1, R Core Team, 2023). Differences in recruit densities between the different shapes (control, open hexagon, closed hexagon, fractal dome, hexagon dome, sizedome, and superdome) after one week, two months, six months, and one year were derived from Wilcoxon tests followed by holm-adjustment for multiple testing. To assess whether the sample sizes were sufficient to detect treatment effects, we conducted a post hoc power analysis using observed means, standard deviations, and sample sizes for each dome type and timepoint. For each pairwise comparison, we calculated Cohen’s d and estimated power using a two-sample approximation. Differences in recruit densities between the different shapes and the natural reef after two months were derived from Wilcoxon tests followed by holm-adjustment for multiple testing. Differences in recruit numbers (total numbers summarized from three replicate recesses of the same depth on one dome) and recruit densities (number of recruits per cm^2^) between the different recess depths (1, 2, 3, and 4 cm) and between the different positions within the recesses (CE: cryptic edge, I: inside, EE: exposed edge, O: outside), split by recess depths (1, 2, 3, and 4 cm) after one week, two months, six months, and one year were derived accordingly. Similarly, differences in coral size and biomass parameters (diameter per recruit, planar surface area per recruit, volume per recruit, total planar surface area per dome, and total volume per dome) on control domes and superdomes at reef 13 and Moku o Loʻe one year were assessed.

The overall impact of flow on coral settlement (recruit density) was assessed using a linear model with flow as a continuous variable, separately for each dome type. Differences in recruit densities between the different flow treatments (no, low, and high) on superdomes and control domes after one week, two months, six months, and one year were derived from Wilcoxon tests followed by holm-adjustment for multiple testing. One dome was excluded from the analysis of the tank experiment (initially in the no flow treatment, later in the low flow treatment) as it was a sizedome, which was included accidentally instead of a superdome.

## 3 Results

### 3.1 Settlement and survival of coral recruits on different module designs

Throughout the experiment, the highest numbers of recruits and recruit densities were found on modules with helix recesses (i.e., fractal dome, hexagon dome, sizedome, and superdome; Figures 2A and S4, Wilcoxon tests, p < 0.05, Tables S2, S3, and S4). Control domes consistently had the lowest number of recruits and recruit densities. One week after coral spawning, remarkably high numbers of recruits were found, with up to 1089 recruits on a single sizedome and 1049 on a single superdome, resulting in densities of ∼1 recruit per cm^2^ (Tables S3 and S4). Recruit densities on modules with helix recesses decreased during the initial post-settlement mortality and stabilized at ∼0.02 recruits per cm^2^ after the first two months. Much lower numbers of recruits were found on modules at Moku o Loʻe, compared to patch reef 13. There, densities increased over time to levels greater than patch reef 13 until they dropped again at the one-year mark. Nonetheless, the overall preference for helix recesses was similar at both sites. Post hoc power analysis confirmed that comparisons between helix-based structures (superdome, sizedome, and hexagon dome) and non-helix designs were well powered (Power > 0.8 in most cases, Table S5). In comparison to natural reef substrate, settlement was significantly higher on all settlement structures, except for the controls after two months (Wilcoxon test, p<0.0001, Table S6).

**Figure 2:**
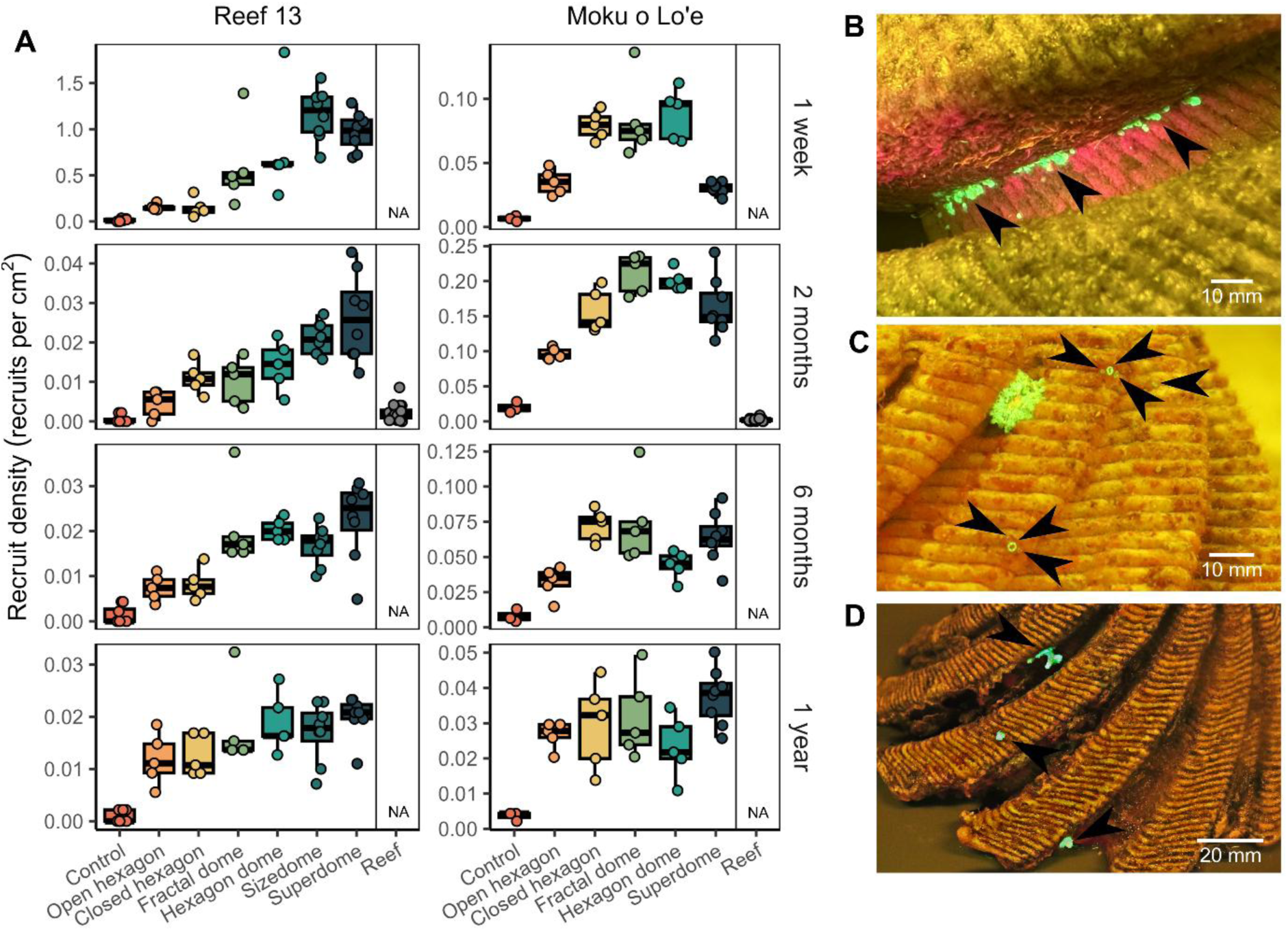
Coral larvae settlement and survival on different settlement modules. (A) Recruit densities (recruits per cm^2^) on the seven different designs (control, open hexagon, closed hexagon, fractal dome, hexagon dome, sizedome, and superdome) at the two study locations: patch reef 13 and Moku o Loʻe after one week, two months, six months, and one year and on natural reef substrate in the Barrier reef in Kāneʻohe Bay after two months. Data are displayed as box-and-whisker plots with raw data points; lines indicate medians, boxes indicate the first and third quartile, and whiskers indicate ± 1.5 inter-quartile range (IQR). (B) *M. capitata* recruits on superdome design from reef 13 after one week. Recruits were mainly found in the corner of the cryptic edge and the inside of helix domes. (C) *M. capitata* recruits on superdome design from reef 13 after two months. (D) *Pocillopora* sp. recruits on superdome design from Moku o Loʻe after one year. Arrows indicate recruits (fluorescent green), highlighted using blue lights and photographed through a yellow filter.

### 3.2 Recess properties of settlement locations of coral recruits

The properties of the recesses where coral recruits were found were analyzed in detail on the high performing superdome modules at reef 13 (Figure 3A, i.e., depth, width, light levels, and positions within the recess). The total number of recruits differed between the recess depths (Figure 3B, Wilcoxon tests p < 0.05, Table S7). Specifically, significantly more recruits settled in recesses with depths of 2, 3, and 4 cm than with a depth of 1 cm (depth measured at the bottom of the module). Initially, most recruits were found in recesses with a depth of 3 cm. Long-term survival was highest in recesses of 4 cm depth. Recruit densities, standardized to the surface area of the respective recesses, showed a similar trend across depths. However, these differences were not statistically significant.(Wilcoxon tests p ≥ 0.05, Table S7).

Coral larvae settled primarily in recess areas with 1.40 ± 0.64 cm width at depths of 1.91 ± 0.82 cm, (Figure 3C, Figure S3B, Table S8). Long-term survival shifted towards larger recess sizes and was highest in recess areas with widths of 1.52 ± 0.63 cm and depths of 1.74 ± 0.79 cm. Relative light levels (based on a light level of ∼200 µmol m^-2^ s^-1^ PAR at the top of the dome) in which coral larvae settled were rather low and coral recruits were found in areas with light levels of 51.8 ± 48.2 µmol m^-2^ s^-1^ PAR (Figure 3D). Long-term survival revealed a larger distribution across light levels and recruits were mainly found in slightly brighter areas with 59.9 ± 48.2 µmol m^-2^ s^-1^ PAR.

Further, the total number of recruits differed between the positions (cryptic edge, inside, exposed edge, and outside; Figure 3E, Wilcoxon tests p < 0.05, Table S9). Independent of recess depth, most recruits were found in the inside corner of the cryptic edge of the recess (Figures 2B and 3E). In contrast, recruits were rarely found on the exposed edge or the outside of the recess. The same patterns were observed for recruit densities, standardized to the area of the respective recess sides (Wilcoxon tests p < 0.05, Table S9).

**Figure 3:**
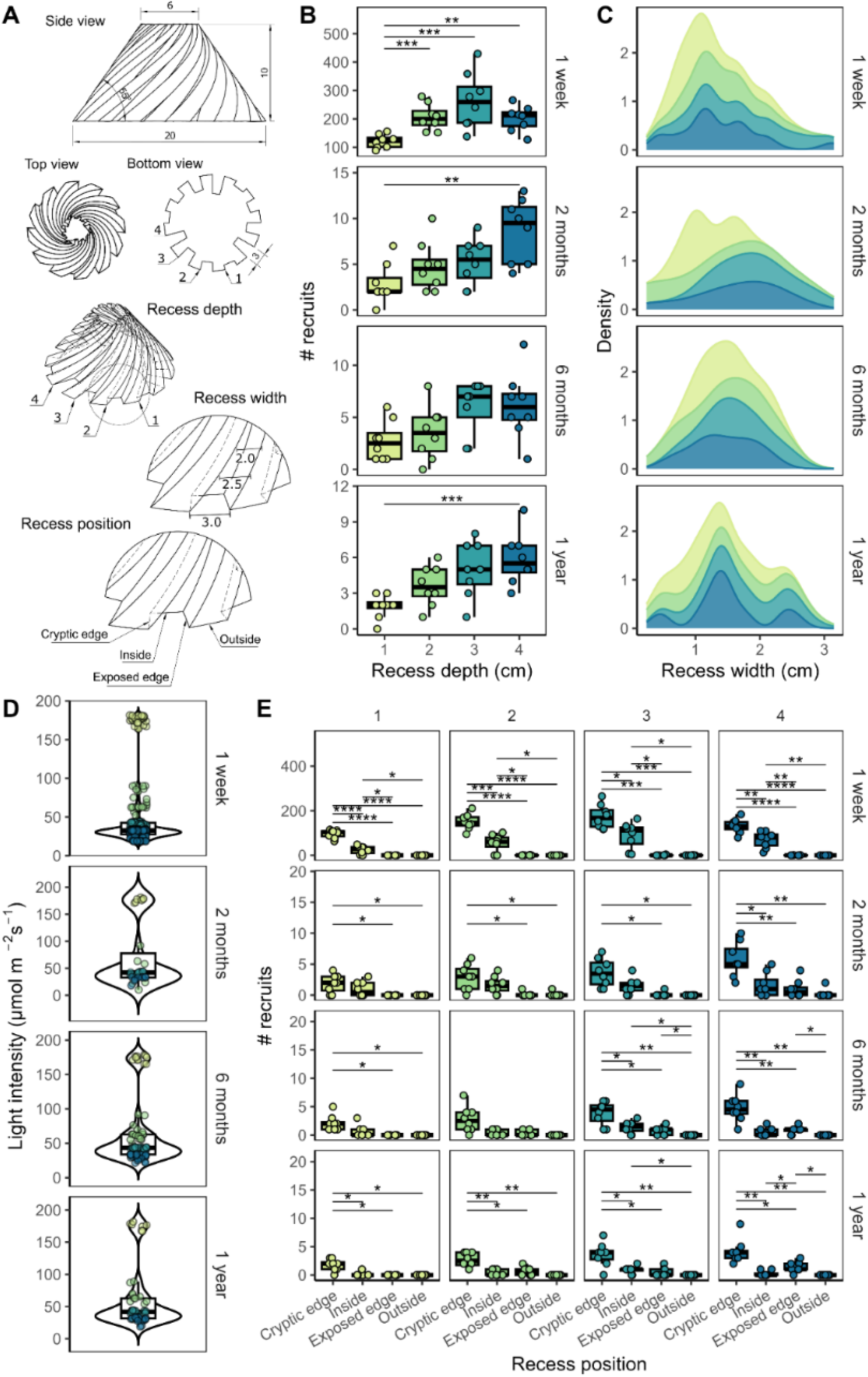
Recess properties of the superdome modules. Recess properties (recess depth, recess width, light levels, and positions within the recess) of the superdome module affecting coral settlement and survival were compared at reef 13 after one week, two months, six months, and one year. Colors indicate the four recess depths (1, 2, 3, and 4 cm depth, measured at the bottom of the module). (A) Detailed drawing of superdome design, basic dimensions of the settlement modules (diameter, height, angles, unit: cm) and the recess properties studied. (B) Total number of recruits in different recess depths (1, 2, 3, and 4 cm). Data are displayed as box-and-whisker plots with raw data points; lines indicate medians, boxes indicate the first and third quartile, and whiskers indicate ± 1.5 IQR. Asterisks indicate levels of statistical significance, derived from Wilcoxon tests followed by holm-adjustment for multiple testing: * p ≤ 0.05, ** p ≤ 0.01, ***p ≤ 0.001. (C) Recess widths where recruits were found on a subset of superdomes. Data are displayed as smoothed density estimates, stacked from the different recess depths. (D) Light levels (µmol m^-2^ s^-1^, PAR) at which recruits were found on a subset of superdomes. Data are displayed as violin plots with inset box-and-whisker plots and raw data points. (E) Total number of recruits at the different positions within the recesses (CE: cryptic edge, I: inside, EE: exposed edge, O: outside), split by recess depths (1, 2, 3, and 4 cm). Data are displayed as box-and-whisker plots. Asterisks indicate levels of statistical significance as above.

### 3.3 Recruit growth and genera found on settlement modules

After one year, coral recruits had grown to a diameter of 3.40 ± 1.91 mm (31−11 mm, min−max here and in the following) on superdome settlement modules at reef 13 and 2.89 ± 1.63 mm (1−12 mm) at Moku o Loʻe (Table S10). This translated to planar surface areas of 11.40 ± 12.4 mm^2^ (0.79− 55 mm^2^) on superdomes at reef 13 and 8.38 ± 11.50 mm^2^ (0.79−104 mm^2^) at Moku o Loʻe. The volume of coral recruits was estimated to be 29.5 ± 47.8 mm^3^ (0.79−220 mm^3^) at reef 13 and 23.7 ± 83.2 mm^3^ (0.79−933 mm^3^) at Moku o Loʻe. Recruit sizes on superdomes did not show a clear trend for being larger or smaller than those on control domes (Wilcoxon tests, Table S11).

We identified coral recruits of the genera *Montipora*, *Pocillopora*, *Leptastrea*, and *Porites* on the settlement modules (Figure 2B−D). However, coral genera could only be assigned to a fraction of the recruits (∼5%, mainly fast-growing *Pocillopora* spp.), as most recruits were too small to be confidently identified. Given this bias, coral genera were not further evaluated. We noted that the initial recruitment was dominated by *Montipora capitata* at reef 13, later followed by additional recruitment by brooding species, such as *Pocillopora* spp. (observational). At Moku o Loʻe, recruitment was dominated by brooding *Pocillopora* spp. and *Leptastrea purpurea*, independent of time.

### 3.4 Coral settlement and survival under different flow regimes

In a separate tank experiment, we compared the effects of three different flow regimes (no, low, and high flow; flow speed of 0, 10, and 15 cm s-1, respectively) on settlement and survival rates of *M. capitata* on coral settlement modules of the superdome and control design. Recruit densities on superdomes increased significantly with higher flow conditions one week after spawning (Linear model, coefficient = 0.019, p = 0.0006, R^2^ = 0.433). Specifically, recruit densities were significantly higher in the ‘medium’ and the ‘high flow’ treatments compared to the ‘no flow’ treatment (Figure 4, Wilcoxon test, p = 0.0410 and p = 0.0110, respectively). Flow had no impact on recruit densities on superdomes after two months or six months. Recruit densities on control domes were not affected initially during settlement but were higher on domes from the ‘high flow’ treatment after two months (Kruskal test, p = 0.0340, Table S12, Figure S5).

**Figure 4:**
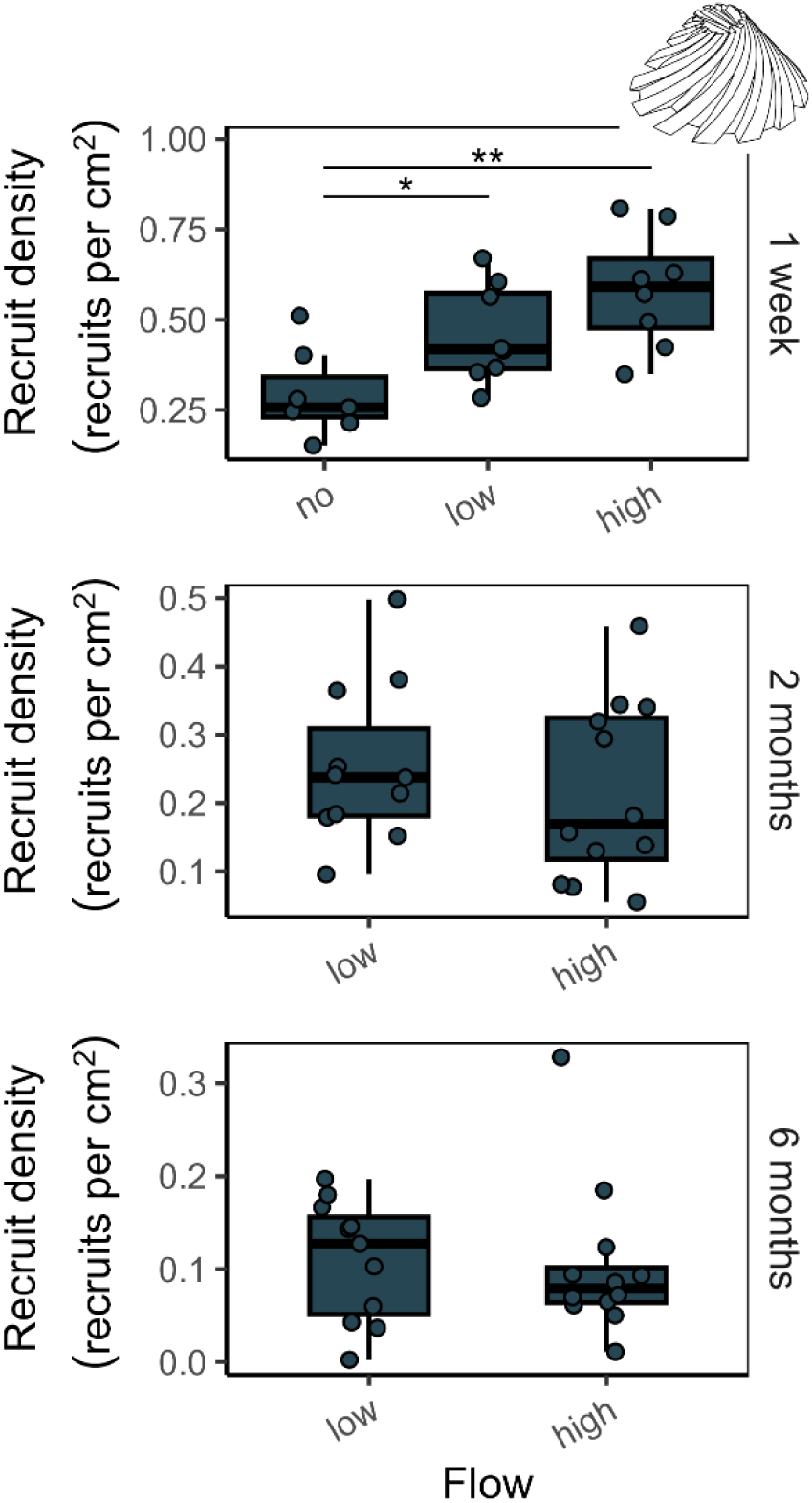
Coral settlement and survival on superdome modules under different flow conditions. Recruit densities (recruits per cm^2^) were assessed on the domes in the experimental tanks after one week, two months, and six months. Settlement was studied under three different flow regimes (no flow, low flow, and high flow) after one week. Survival was studied under two different flow regimes (low flow and high flow) after two months and six months. Data are displayed as box-and-whisker plots with raw data points; lines indicate medians, boxes indicate the first and third quartile, and whiskers indicate ± 1.5 IQR. Significant differences are marked with asterisks, defined as p < 0.001 (***), p < 0.01 (**), and p < 0.05 (*), and derived from Wilcoxon tests followed by holm-adjustment for multiple testing (one week) and Kruskal test (two months and six months).

## 4 Discussion

### 4.1 Helix designs promote settlement and survival of coral recruits

We present a set of optimized recess designs suitable for the implementation into artificial reefs and breakwater systems. The designs increase coral settlement by 80-fold, and post-settlement survival by 20−50-fold, when compared with controls. The identified dimensions are comparable to the micro-scale complexity of benthic substrata (in the scale of 1–10 mm) in reefs (Brandl et al., 2014; Doropoulos et al., 2016; Edmunds et al., 2004), but reveal dramatically higher densities of recruits (increased by ∼70-fold, compared to the natural recruitment measured after two months, on average 0.003 recruits per cm^2^). The numbers of recruits we found at the reef are comparable to what has been previously found on natural substrates in Hawaiʻi (1−2 recruits per m^2^, Brown, 2004). Similarly, settlement on control domes was much lower than on the other coral settlement structures, indicating that the flat surface presents a less attractive settlement substrate despite the same micro-scale topography produced by the 3D printing process, which is more exposed to external impacts, such as grazing and competition with benthic algae (Doropoulos et al., 2016). Modules with holes and pockets resulting from opposing helix patterns (open and closed hexagon) revealed lower settlement than modules with recesses, yet higher long-term survival rates. Reasons for these differences might be the higher light levels in these spaces (compare Figure S3), and the lower presence of sessile invertebrates and algae competing for space with the coral recruits (Chadwick and Morrow, 2011). Also, holes may allow for better water circulation, which is crucial for gas exchange, nutrients, and food supply (Mass et al., 2010; Sebens et al., 1997), while recesses may have more restricted water flow. Thus, future design iterations might incorporate recesses providing higher light levels and increased flow.

Although the overall settlement patterns between the designs were largely consistent over time, with helix designs outperforming controls, strong differences in the total numbers of recruits were observed. The initially extremely high recruit numbers (up to > 1000 recruits per settlement module) were substantially reduced through post-settlement mortality. These early-life bottlenecks are inherently part of the life history strategy of spawning corals, following a Type III survivorship curve where mortality is highest immediately after settlement (Gosselin and Qian, 1997; Martinez and Abelson, 2013; Nozawa and Harrison, 2008). Future developments can include mitigation methods improving post-settlement survivorship. For example, by pre-seeding settlement modules with CCA to prevent the growth of stronger competitors such as fleshy algae (Vermeij et al., 2011).

Additionally, we observed differences between the two field study locations that converged over time. At patch reef 13, the total number of recruits per module was much higher than at Moku o Loʻe. There, settlement modules were exposed to a medium flow environment located at the channel of Kāneʻohe Bay, promoting a high availability of coral larvae in the water column during spawning. The initial recruitment was dominated by *Montipora capitata*, followed later by additional recruitment by monthly brooding species, such as *Pocillopora* spp. This presents an example of a high seeding event after the mass spawning of *M. capitata*, leading to an initial high recruitment followed by high post-settlement mortality. Despite these dynamics, the relative trends in settlement and survival across the different module shapes remained stable over time. In contrast, initial recruitment at Moku o Loʻe was comparatively low. There, settlement modules were exposed to a shallower water environment, with higher daily temperature fluctuations (compare Figure S6) and lower water flow. The recruitment was dominated by more hardy, brooding coral species, such as *Pocillopora* spp. and *Leptastrea purpurea* (Bahr et al., 2016), which release larvae in repetitive spawning events during the summer months. This presents an example of a successive seeding event, with low initial but continuous recruitment over time and fluctuations in mortality. However, similar to the observation at reef 13, while variability was observed over time, the overall trends of recruitment and survival on different module shapes remained largely consistent. Although the two sites had different environmental conditions and initial recruitment patterns, the performance rankings of the module designs were similar. This indicates that the structural features identified consistently enhance coral recruitment across diverse reef environments and coral communities. Notably, final coral recruitment on helix modules converged across sites despite varying larval availability. This indicates that structures with helix recesses may perform reliably across different reef environments, regardless of local larval supply, and can be widely used.

### 4.2 Recess properties affect settlement and survival of coral recruits

Positions at which recruits have settled within the recess were measured to identify preferred recess dimensions. We found that recesses with widths of approximately 15 mm and depths of around 17 mm were most conducive to coral settlement at reef 13. Light levels in these areas were comparably low ∼56 µmol m^-2^ s^-1^, corresponding to ∼28% of the incident light on top of the module. While this provides useful information on relative light gradients between surface and crevice habitats, it is important to note that these absolute values are only approximations and the actual light environment in the field may differ due to diel light variability. Consequently, these values are proxies for structural comparison, not precise representations of *in situ* light exposure or daily light integral (DLI). Nonetheless, the observed light attenuation patterns align with known coral larval preferences for microhabitats. Coral larvae prefer moderate light exposure, avoiding overly shaded or extremely bright locations. Similar recess sizes and low light levels have been observed to be beneficial for the survival of coral recruits (Doropoulos et al., 2016; Rahnke et al., 2022; Ramsby et al., 2024). However, the response of recruits to light intensity can be mixed, with other studies showing evidence of faster growth or higher survival under high light intensity (Hancock et al., 2021; Koch et al., 2022). Over time, initial settlement and survival only differed in the mm scale, shifting from slightly smaller recess areas during initial settlement towards larger recess sizes for long-term survival. Similarly, coral recruits were found to have settled in areas with lower light levels of ∼ 50 µmol m^-2^ s^-1^, but survival was higher under higher light levels of ∼60 µmol m^-2^ s^-1^. This is likely due to the presence of competitors such as sessile invertebrates (i.e., sponges or oysters) in larger and darker areas of the recesses (Chadwick and Morrow, 2011). Overall, a great variety of other benthic organisms, such as sponges, tunicates, and oysters, colonized the surfaces. Sponges were especially found competing with coral recruits inside crevices, but these observations are pending further analyses. These interactions did not impair module integrity and all modules remained structurally stable over the one-year deployment, with no signs of degradation or sediment accumulation.

The settlement preferences of coral larvae are a trade-off between light-capture, potential to be overgrown by competitive algae in early life stages and the avoidance of grazing (Doropoulos et al., 2016). Thus, a successful strategy is to settle in recesses near underside edges where they are protected from predation, grazing and strong physical impacts but still receive sufficient light to grow, and from which they then grow out into brighter, more open spaces (Cruz and Harrison, 2017). This corresponds well with the positions at which we found recruits inside the recess: independent of the depth of the recess, most recruits were found in the inside corner of the cryptic edge of the recesses, which provide shelter and lower exposure. In contrast, recruits were rarely found on the exposed edge and the outside of the recess, where they might be more exposed to predation, high light, and competition with fleshy algae.

### 4.3 Coral settlement on helix modules is increased under higher flow

We found that settlement on superdomes was significantly increased under higher flow conditions. This suggests that settlement inside the recesses can be strongly influenced by hydrodynamics. Hydrodynamics play an important role in the distribution and settlement of coral larvae, and might have potentially trapped coral larvae through the formation of turbulent flow and micro-eddies (Gysbers et al., 2024; Reidenbach et al., 2009). Future hydrodynamic measurements and models are needed to test this hypothesis. The millimeter-scale topography of ridges, formed through the layered printing of the clay, might have additionally aided coral settlement in the sheltered areas of the complex modules, by increasing areas of recirculation, directing larvae toward the structure and increasing the window of time for the larvae to settle (Levenstein et al., 2022a). However, given the low levels of settlement on control modules and the outside surfaces of other modules, the small-scale topography alone did not greatly enhance settlement and may only be beneficial in the presence of meso-scale features such as recesses. Regarding survival, flow had no impact on recruits on superdomes, where recruits are rather sheltered inside the recesses. In contrast, on control domes, survival was higher under higher flow, which is in accordance with previous studies that found positive impacts of water flow for the survival of coral recruits (Hancock et al., 2021; Rahnke et al., 2022). On the control domes, recruits are more directly exposed to flow conditions, and the positive effects of increased flow on coral physiology, such as gas exchange, nutrient supply, and food availability, become more evident (Mass et al., 2010; Sebens et al., 1997). These results suggest that flow has an important impact on settlement on the meso- to micro-scale (m to mm range). Consequently, sites with moderate to high water flow may be particularly suitable for maximizing the effectiveness of these designs.

### 4.4 Limitations, future directions, and applications

We demonstrate that artificial structures with helix recesses can dramatically promote coral settlement and survival of common coral species at two reef sites in Kāneʻohe Bay, Hawaiʻi. This success is probably due to the sheltered microhabitats formed by the helix recesses, advantageous hydrodynamic conditions, and the increased functional settlement surface area in helix designs, all of which improve larval retention and settlement chances. The recruits found mainly belonged to *Montipora* spp. and *Pocillopora* spp., which are dominant reef-builders in the reefs studied. It remains to be tested how the structures perform in other reefs around the world, where other spawning species, such as *Acropora* spp., dominate the reefs. While strong trends in settlement and survival were observed across designs, survival data combined all coral recruits regardless of genus. *Montipora* sp. (spawner) and *Pocillopora* sp. (brooder) differ in reproductive strategies and survival patterns, but recruits could not be reliably distinguished due to their small size and high densities (up to 900 per module). Continued post-spawning recruitment might have obscured early mortality patterns. Future work should include species-level identification (e.g., genetic or through bleaching and corallite analysis) and individual tracking to resolve genus-specific dynamics. Further, in this study, we tested a rather narrow flow range (i.e., 0, 10, and 15 cm s^-1^), corresponding to average conditions in coral reefs (Roik et al., 2016). Higher flow speeds, occurring in high-flow environments of coral reefs, with peak speeds of 30−40 cm s^-1^ (Lentz et al., 2017), might have a negative impact on settlement (Koehl, 2007), resulting in a hump-shape relationship between flow and coral recruitment, and need to be further explored. While our structures provided especially high initial settlement rates, essentially trapping larvae from the water column, recruit numbers were substantially reduced through post-settlement mortality. Future design iterations and biological interventions might target post-settlement survival, for example, by incorporating recesses providing higher light and flow levels or interventions decreasing benthic competitors. The integration and application of the findings is straightforward and can greatly enhance coral restoration efforts. Helix recess designs can be integrated into local restoration modules, large-scale coastal protection structures, and other marine infrastructure projects with nature-positive aspirations or mandates. This structure-based approach passively enhances coral recruitment and survival, and reduces the need for costly and labor-intensive coral transplantation. The increased efficiency and scalability of this approach could have significant long-term ecological and financial benefits, including increased return on restoration investment and improvements of ecosystem services, such as shoreline protection, fisheries, and tourism. To encourage broader adoption, we advocate for establishing a dedicated fund for modular coral reef restoration efforts, specifically focusing on coral larvae. We aim to develop design standards for integrating helix features into reef structures in the future, to better align engineering methods with ecological restoration. These measures will provide the basis for more effective, scalable, and self-sustaining artificial or hybrid reefs that enhance biodiversity and strengthen coastal resilience.

## Data and materials availability

Data and scripts are available from GitHub https://github.com/JessiReichert/Helix_coral_settlement.

## Acknowledgements

We thank Marion Chapeau, Guan-Yan Chen, Jon Ehrenberg, Peter Felicijan, and Marina Rottmüller for their support during the experiments. Coral larvae were collected under the Hawaiʻi Institute of Marine Biology Special Activity Permit 2024-45. We thank the students of the 2023 MBIO640 class at the University of Hawaiʻi for helping collect the reef recruit data. This work was funded by the Defense Advanced Research Projects Agency (Contract No. HR001122C0134), the National Science Foundation (1948946), and the HIMB Director’s Innovation Fund.

## Author contributions

Conceptualization: JR, HJ, NMDS, and JSM

Methodology: JR, HJ, NMDS, and JSM

Investigation: JR, HJ, CD, JRH, CH, AN, and LHR

Visualization: JR

Supervision: JSM

Writing—original draft: JR

Writing—review & editing: JR, HJ, NMDS, and JSM

## Use of generative AI

During the preparation of this work the large language model Gemini, (Google AI) and ChatGPT (OpenAI) were used to improve the language of the manuscript. The authors thoroughly reviewed all content and take full responsibility for the content of the published article.

