## Supplementary Materials for "Helix recesses boost coral larvae settlement and survival"

**This PDF file includes**

Figs. S1 to S6
Tables S1 to S11

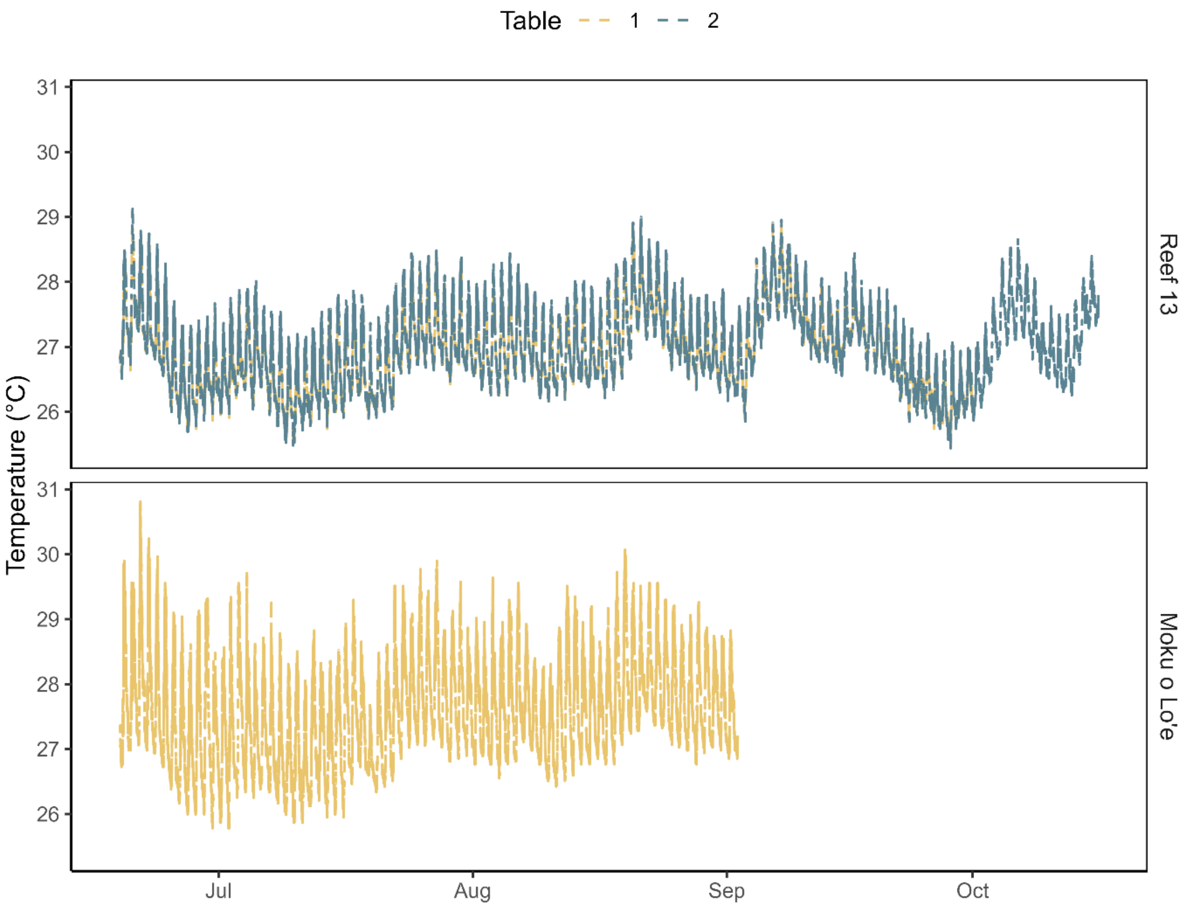


**Figure S1.**
Temperature (°C) in the field locations reef 13 and Moku o Lo‘e. Temperatures were recorded using HOBO data loggers with intervals of 5 min. The loggers stopped working in September and October 2023 and did not record further despite reinstallation.

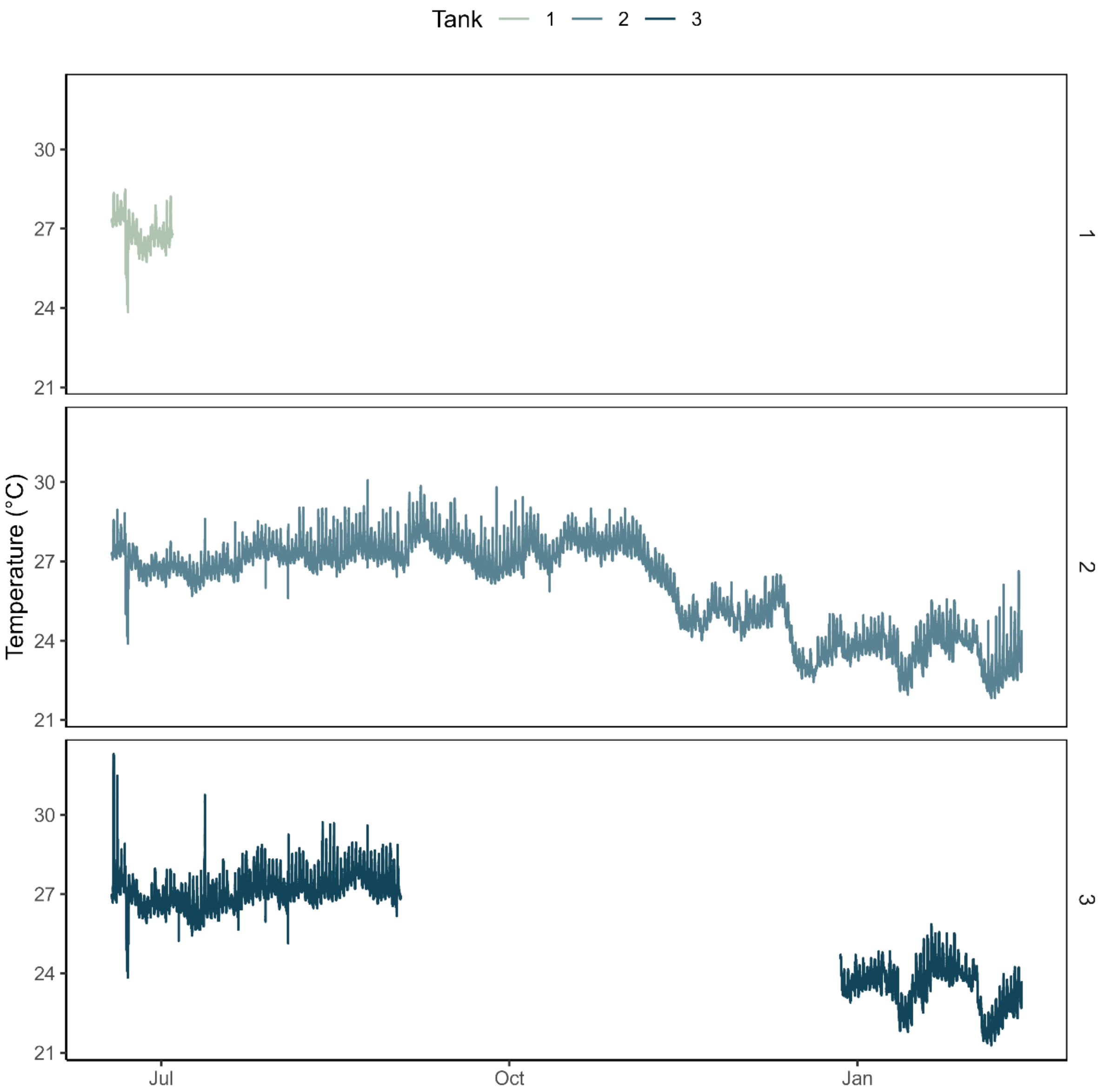


**Figure S2.**
Temperature (°C) in the tanks. Temperatures were recorded using HOBO data loggers with intervals of 5 min. Tank 1 was only used for the seeding and during the initial counting of the coral larvae. The logger in tank 3 stopped working in September and was reinstalled in December 2023.

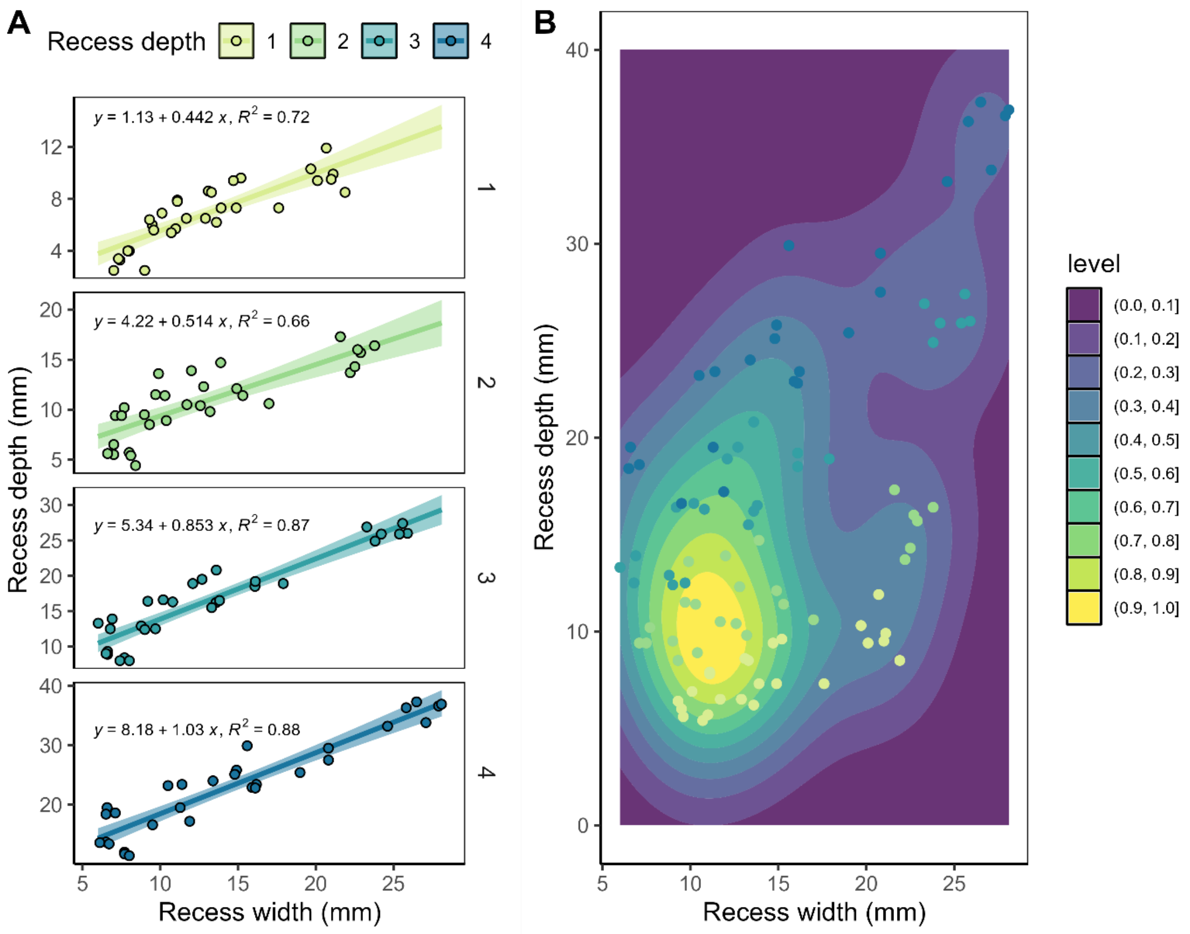


**Figure S3.**

Recess width-depth space on superdomes. (A) Recess width and depth assessed at five positions (base, 2 cm height, 4 cm height, 6 cm height, and 8 cm height) for each recess depth (1, 2, 3, and 4 cm) on two superdomes. Linear models (solid lines) are derived from raw data (points) and given with standard deviation (light-colored area), equation and $R^2$. (B) Recess width-depth space present on the superdome. Data are displayed as raw data (points) and kernel density estimations with bright values (yellow) indicating high recruit densities and dark values (purple) indicating low recruit densities.

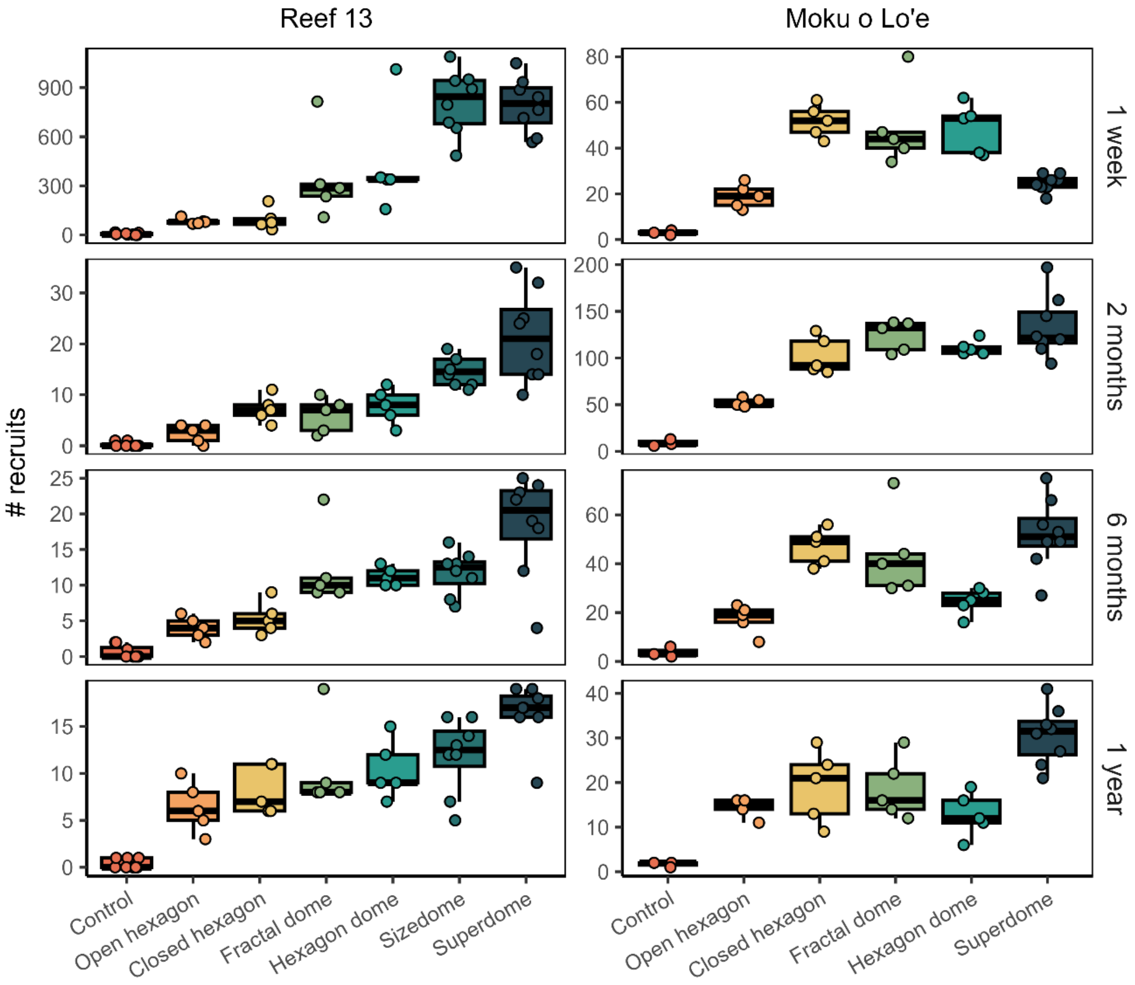


**Figure S4.**

Coral larvae settlement and survival on different settlement modules. Recruit numbers (total numbers per dome) on the seven different designs (control, open hexagon, closed hexagon, fractal dome, hexagon dome, sizedome, and superdome) at the two study locations reef 13 and Moku o Lo‘e after one week, two months, six months, and one year. Data are displayed as box-and-whisker plots with raw data points; lines indicate medians, boxes indicate the first and third quartile, and whiskers indicate ± 1.5 IQR.

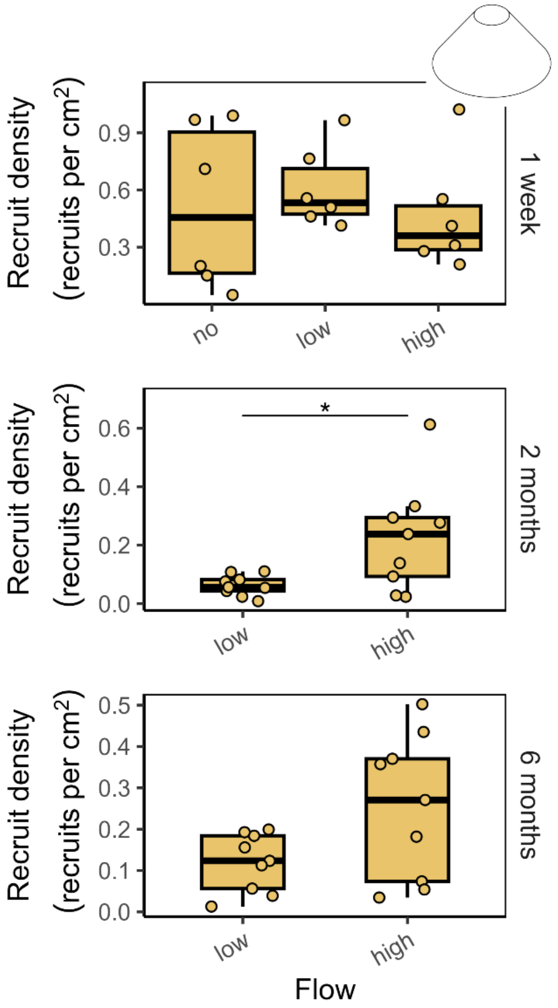


**Figure S5.**

Coral larvae recruitment and survival on control dome modules under different flow conditions. Recruit densities (recruits per $cm^2$) were assessed on the domes in the experimental tanks after one week, two months, and six months. Settlement was studied under three different flow regimes (no flow, low flow, and high flow) after one week. Survival was studied under two different flow regimes (low flow and high flow) after two months and six months. Data are displayed as box-and-whisker plots with raw data points; lines indicate medians, boxes indicate the first and third quartile, and whiskers indicate ± 1.5 IQR. Significant differences are marked with asterisks, defined as $p < 0.001$ (***), $p < 0.01$ (**), and $p < 0.05$ (*), and derived from Wilcoxon tests followed by holm-adjustment for multiple testing (one week) and Kruskal test (two months and six months).

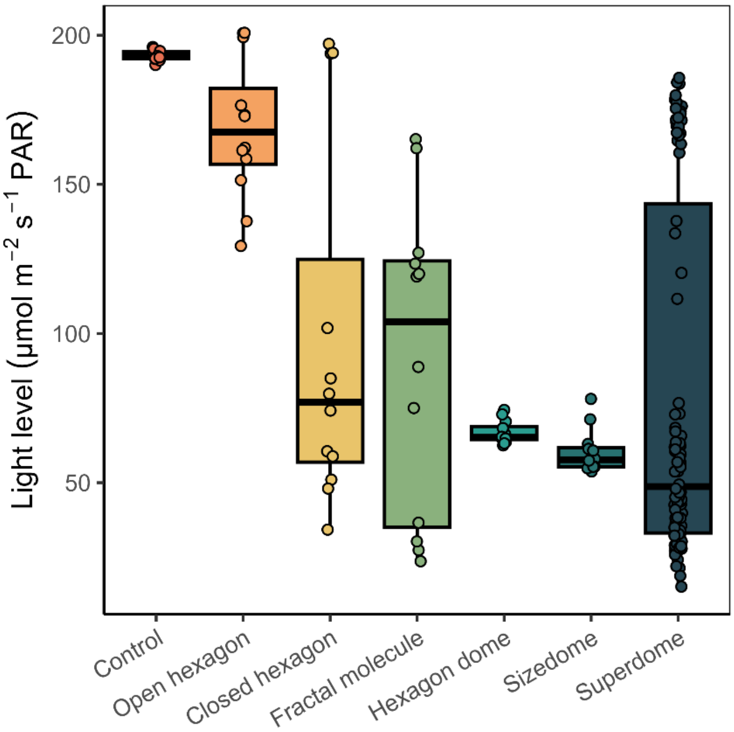


**Figure S6.**
Light levels on settlement modules. Light levels were measured on the seven different settlement modules (control, open hexagon, closed hexagon, fractal dome, hexagon dome, sizedome, and superdome) in the settlement habitats at four positions inside the recesses (the base of the dome, 2 cm height, 4 cm height, and 6 cm height) or in the holes, respectively. Data are displayed as box-and-whisker plots with raw data points; lines indicate medians, boxes indicate the first and third quartile, and whiskers indicate ± 1.5 IQR.

**Table S1.**

Number of replicates for the seven designs at the two field locations.

| Design | Reef 13 | Moku o Lo‘e |
|---|---|---|
| Control | 8 | 3 |
| Open hexagon | 5 | 5 |
| Closed hexagon | 5 | 5 |
| Fractal dome | 5 | 5 |
| Hexagon dome | 5 | 5 |
| Sizedome | 8 | 0 |
| Superdome | 8 | 8 |

**Table S2.**

Differences in recruit densities (recruits per $cm^2$) between the different shapes (control, open hexagon, closed hexagon, fractal dome, hexagon dome, sizedome, and superdome) after one week, two months, six months, and one year. Results are derived from Wilcoxon tests followed by holm-adjustment for multiple testing. Number of replicates per test group (n1 and n2), test statistic ($chi^2$), and p-value are given with bold values indicating significance ($p < 0.05$).

| Timepoint | Contrast | | | n1 | n2 | Statistic | p |
|---|---|---|---|---|---|---|---|
| 1 week | Control dome | - | Open hexagon | 11 | 10 | 4 | **0.0052** |
| | Control dome | - | Closed hexagon | 11 | 10 | 0 | **0.0022** |
| | Control dome | - | Fractal dome | 11 | 10 | 0 | **0.0022** |
| | Control dome | - | Hexagon dome | 11 | 10 | 0 | **0.0022** |
| | Control dome | - | Sizedome | 11 | 8 | 0 | **0.0048** |
| | Control dome | - | Superdome | 11 | 16 | 7 | **0.0013** |
| | Open hexagon | - | Closed hexagon | 10 | 10 | 42 | 1 |
| | Open hexagon | - | Fractal dome | 10 | 10 | 24 | 0.5200 |
| | Open hexagon | - | Hexagon dome | 10 | 10 | 25 | 0.5760 |
| | Open hexagon | - | Sizedome | 10 | 8 | 0 | **0.0010** |
| | Open hexagon | - | Superdome | 10 | 16 | 64 | 1 |
| | Closed hexagon | - | Fractal dome | 10 | 10 | 33 | 1 |
| | Closed hexagon | - | Hexagon dome | 10 | 10 | 28 | 1 |
| | Closed hexagon | - | Sizedome | 10 | 8 | 0 | **0.0010** |
| | Closed hexagon | - | Superdome | 10 | 16 | 80 | 1 |
| | Fractal dome | - | Hexagon dome | 10 | 10 | 42 | 1 |
| | Fractal dome | - | Sizedome | 10 | 8 | 7 | **0.0260** |
| | Fractal dome | - | Superdome | 10 | 16 | 88 | 1 |
| | Hexagon dome | - | Sizedome | 10 | 8 | 8 | 0.0600 |
| | Hexagon dome | - | Superdome | 10 | 16 | 88 | 1 |
| | Sizedome | - | Superdome | 8 | 16 | 108 | 0.0880 |
| 2 months | Control dome | - | Open hexagon | 11 | 10 | 22 | 0.3000 |
| | Control dome | - | Closed hexagon | 11 | 10 | 14 | 0.0720 |
| | Control dome | - | Fractal dome | 11 | 10 | 13 | 0.0570 |
| | Control dome | - | Hexagon dome | 11 | 10 | 10 | 0.0400 |
| | Control dome | - | Sizedome | 11 | 8 | 11 | 0.1020 |
| | Control dome | - | Superdome | 11 | 16 | 8 | 0.0017 |
| | Open hexagon | - | Closed hexagon | 10 | 10 | 27 | 1 |
| | Open hexagon | - | Fractal dome | 10 | 10 | 31 | 1 |
| | Open hexagon | - | Hexagon dome | 10 | 10 | 28 | 1 |
| | Open hexagon | - | Sizedome | 10 | 8 | 40 | 1 |
| | Open hexagon | - | Superdome | 10 | 16 | 40 | 0.5180 |
| | Closed hexagon | - | Fractal dome | 10 | 10 | 41 | 1 |
| | Closed hexagon | - | Hexagon dome | 10 | 10 | 36 | 1 |
| | Closed hexagon | - | Sizedome | 10 | 8 | 41 | 1 |
| | Closed hexagon | - | Superdome | 10 | 16 | 59 | 1 |
| | Fractal dome | - | Hexagon dome | 10 | 10 | 47 | 1 |
| | Fractal dome | - | Sizedome | 10 | 8 | 41 | 1 |
| | Fractal dome | - | Superdome | 10 | 16 | 74 | 1 |
| | Hexagon dome | - | Sizedome | 10 | 8 | 48 | 1 |
| | Hexagon dome | - | Superdome | 10 | 16 | 79 | 1 |
| | Sizedome | - | Superdome | 8 | 16 | 21 | 0.1440 |
| 6 months | Control dome | - | Open hexagon | 11 | 10 | 10 | 0.0320 |
| | Control dome | - | Closed hexagon | 11 | 10 | 6 | 0.0098 |
| | Control dome | - | Fractal dome | 11 | 10 | 0 | 0.0022 |
| | Control dome | - | Hexagon dome | 11 | 10 | 0 | 0.0022 |
| | Control dome | - | Sizedome | 11 | 8 | 2 | 0.0095 |
| | Control dome | - | Superdome | 11 | 16 | 2 | 0.0005 |
| | Open hexagon | - | Closed hexagon | 10 | 10 | 36 | 1 |
| | Open hexagon | - | Fractal dome | 10 | 10 | 18 | 0.2210 |
| | Open hexagon | - | Hexagon dome | 10 | 10 | 25 | 0.7040 |
| | Open hexagon | - | Sizedome | 10 | 8 | 35 | 1 |
| | Open hexagon | - | Superdome | 10 | 16 | 40 | 0.4440 |
| | Closed hexagon | - | Fractal dome | 10 | 10 | 41 | 1 |

| | | | | | | | |
|---|---|---|---|---|---|---|---|
| | Closed hexagon | - | Hexagon dome | 10 | 10 | 50 | 1 |
| | Closed hexagon | - | Sizedome | 10 | 8 | 42 | 1 |
| | Closed hexagon | - | Superdome | 10 | 16 | 69 | 1 |
| | Fractal dome | - | Hexagon dome | 10 | 10 | 56 | 1 |
| | Fractal dome | - | Sizedome | 10 | 8 | 61 | 0.7040 |
| | Fractal dome | - | Superdome | 10 | 16 | 79 | 1 |
| | Hexagon dome | - | Sizedome | 10 | 8 | 69 | 0.1540 |
| | Hexagon dome | - | Superdome | 10 | 16 | 55 | 1 |
| | Sizedome | - | Superdome | 8 | 16 | 15 | 0.0450 |
| 1 year | Control dome | - | Open hexagon | 11 | 10 | 0 | 0.0021 |
| | Control dome | - | Closed hexagon | 11 | 10 | 0 | 0.0021 |
| | Control dome | - | Fractal dome | 11 | 10 | 0 | 0.0021 |
| | Control dome | - | Hexagon dome | 11 | 10 | 0 | 0.0021 |
| | Control dome | - | Sizedome | 11 | 8 | 0 | 0.0043 |
| | Control dome | - | Superdome | 11 | 16 | 0 | 0.0003 |
| | Open hexagon | - | Closed hexagon | 10 | 10 | 50 | 1 |
| | Open hexagon | - | Fractal dome | 10 | 10 | 38 | 1 |
| | Open hexagon | - | Hexagon dome | 10 | 10 | 45 | 1 |
| | Open hexagon | - | Sizedome | 10 | 8 | 47 | 1 |
| | Open hexagon | - | Superdome | 10 | 16 | 44 | 0.7930 |
| | Closed hexagon | - | Fractal dome | 10 | 10 | 39 | 1 |
| | Closed hexagon | - | Hexagon dome | 10 | 10 | 44 | 1 |
| | Closed hexagon | - | Sizedome | 10 | 8 | 39 | 1 |
| | Closed hexagon | - | Superdome | 10 | 16 | 42 | 0.6720 |
| | Fractal dome | - | Hexagon dome | 10 | 10 | 57 | 1 |
| | Fractal dome | - | Sizedome | 10 | 8 | 54 | 1 |
| | Fractal dome | - | Superdome | 10 | 16 | 60 | 1 |
| | Hexagon dome | - | Sizedome | 10 | 8 | 49 | 1 |
| | Hexagon dome | - | Superdome | 10 | 16 | 45 | 1 |
| | Sizedome | - | Superdome | 8 | 16 | 18 | 0.0750 |

**Table S3.**

Summary statistics of recruit numbers on modules of different shapes (control, open hexagon, closed hexagon, fractal dome, hexagon dome, sizedome, and superdome) after one week, two months, six months and one year. The number of replicates (n), minimum (min) and maximum (max) values are given with mean and standard deviation (sd).

| Timepoint | Location | Dome type | n | min | max | mean | sd |
|---|---|---|---|---|---|---|---|
| 1 week | Reef 13 | Control | 8 | 0 | 16 | 6.13 | 5.96 |
| | | Open hexagon | 5 | 69 | 113 | 83.40 | 17.50 |
| | | Closed hexagon | 5 | 34 | 206 | 96.20 | 65.80 |
| | | Fractal dome | 5 | 108 | 815 | 351.20 | 270.81 |
| | | Control hexagon | 5 | 158 | 1012 | 440.40 | 329.57 |
| | | Sizedome | 8 | 485 | 1089 | 811.88 | 195.25 |
| | | Superdome | 8 | 567 | 1049 | 793.50 | 167.27 |
| | Moku o Loʻe | Control | 3 | 2 | 4 | 3.00 | 1.00 |
| | | Open hexagon | 5 | 13 | 26 | 19.00 | 5.24 |
| | | Closed hexagon | 5 | 43 | 61 | 51.80 | 7.12 |
| | | Fractal dome | 5 | 34 | 80 | 49.00 | 18.00 |
| | | Control hexagon | 5 | 37 | 62 | 48.80 | 10.90 |
| | | Superdome | 8 | 18 | 29 | 24.75 | 3.62 |
| 2 months | Reef 13 | Control | 8 | 0 | 1 | 0.25 | 0.46 |
| | | Open hexagon | 5 | 0 | 4 | 2.40 | 1.82 |
| | | Closed hexagon | 5 | 4 | 11 | 7.20 | 2.59 |
| | | Fractal dome | 5 | 2 | 10 | 6.00 | 3.39 |
| | | Control hexagon | 5 | 3 | 12 | 7.80 | 3.49 |
| | | Sizedome | 8 | 11 | 19 | 14.63 | 2.88 |
| | | Superdome | 8 | 10 | 35 | 21.50 | 9.01 |
| | Moku o Loʻe | Control | 3 | 6 | 13 | 9.00 | 3.61 |
| | | Open hexagon | 5 | 48 | 58 | 52.00 | 4.30 |
| | | Closed hexagon | 5 | 85 | 129 | 102.40 | 19.81 |
| | | Fractal dome | 5 | 104 | 138 | 124.00 | 16.23 |
| | | Control hexagon | 5 | 105 | 124 | 111.00 | 7.84 |
| | | Superdome | 8 | 94 | 197 | 133.63 | 33.00 |
| 6 months | Reef 13 | Control | 8 | 0 | 2 | 0.63 | 0.92 |
| | | Open hexagon | 5 | 2 | 6 | 4.00 | 1.58 |
| | | Closed hexagon | 5 | 3 | 9 | 5.40 | 2.30 |
| | | Fractal dome | 5 | 9 | 22 | 12.20 | 5.54 |
| | | Control hexagon | 5 | 10 | 13 | 11.20 | 1.30 |
| | | Sizedome | 8 | 7 | 16 | 11.75 | 3.01 |
| | | Superdome | 8 | 4 | 25 | 18.38 | 7.15 |
| | Moku o Loʻe | Control | 3 | 2 | 6 | 3.67 | 2.08 |
| | | Open hexagon | 5 | 8 | 23 | 17.40 | 5.86 |
| | | Closed hexagon | 5 | 38 | 56 | 47.00 | 7.38 |
| | | Fractal dome | 5 | 30 | 73 | 43.60 | 17.47 |
| | | Control hexagon | 5 | 16 | 30 | 24.40 | 5.41 |
| | | Superdome | 8 | 27 | 75 | 52.13 | 14.57 |
| 1 year | Reef 13 | Control | 8 | 0 | 1 | 0.38 | 0.52 |
| | | Open hexagon | 5 | 3 | 10 | 6.40 | 2.70 |
| | | Closed hexagon | 5 | 6 | 11 | 8.20 | 2.59 |
| | | Fractal dome | 5 | 8 | 19 | 10.40 | 4.83 |
| | | Control hexagon | 5 | 7 | 15 | 10.40 | 3.13 |
| | | Sizedome | 8 | 5 | 16 | 11.88 | 3.98 |
| | | Superdome | 8 | 9 | 19 | 16.38 | 3.20 |
| | Moku o Loʻe | Control | 3 | 1 | 2 | 1.67 | 0.58 |
| | | Open hexagon | 5 | 11 | 16 | 14.40 | 2.07 |
| | | Closed hexagon | 5 | 9 | 29 | 19.20 | 8.14 |
| | | Fractal dome | 5 | 12 | 29 | 18.60 | 6.91 |
| | | Control hexagon | 5 | 6 | 19 | 12.80 | 4.97 |
| | | Superdome | 8 | 21 | 41 | 30.63 | 6.48 |

**Table S4.**

Summary statistics of recruit densities (numbers per $cm^2$) on the different shapes (control, open hexagon, closed hexagon, fractal dome, hexagon dome, sizedome, and superdome) after one week, two months, six months and one year. The number of replicates (n), minimum (min) and maximum (max) values are given with mean and standard deviation (sd).

| Timepoint | Location | Dome type | n | min | max | mean | sd |
|---|---|---|---|---|---|---|---|
| 1 week | Reef 13 | Control | 8 | 0.000 | 0.035 | 0.013 | 0.013 |
| | | Open hexagon | 5 | 0.128 | 0.209 | 0.154 | 0.032 |
| | | Closed hexagon | 5 | 0.052 | 0.316 | 0.148 | 0.101 |
| | | Fractal dome | 5 | 0.184 | 1.389 | 0.599 | 0.462 |
| | | Control hexagon | 5 | 0.286 | 1.834 | 0.798 | 0.597 |
| | | Sizedome | 8 | 0.693 | 1.555 | 1.159 | 0.279 |
| | | Superdome | 8 | 0.694 | 1.284 | 0.972 | 0.205 |
| | Moku o Loʻe | Control | 3 | 0.004 | 0.009 | 0.006 | 0.002 |
| | | Open hexagon | 5 | 0.024 | 0.048 | 0.035 | 0.010 |
| | | Closed hexagon | 5 | 0.066 | 0.094 | 0.079 | 0.011 |
| | | Fractal dome | 5 | 0.058 | 0.136 | 0.084 | 0.031 |
| | | Control hexagon | 5 | 0.067 | 0.112 | 0.088 | 0.020 |
| | | Superdome | 8 | 0.022 | 0.036 | 0.030 | 0.004 |
| 2 months | Reef 13 | Control | 8 | 0 | 0.002 | 0.001 | 0.001 |
| | | Open hexagon | 5 | 0 | 0.007 | 0.004 | 0.003 |
| | | Closed hexagon | 5 | 0.006 | 0.017 | 0.011 | 0.004 |
| | | Fractal dome | 5 | 0.003 | 0.017 | 0.010 | 0.006 |
| | | Control hexagon | 5 | 0.005 | 0.022 | 0.014 | 0.006 |
| | | Sizedome | 8 | 0.016 | 0.027 | 0.021 | 0.004 |
| | | Superdome | 8 | 0.012 | 0.043 | 0.026 | 0.011 |
| | Moku o Loʻe | Control | 3 | 0.013 | 0.028 | 0.019 | 0.008 |
| | | Open hexagon | 5 | 0.089 | 0.107 | 0.096 | 0.008 |
| | | Closed hexagon | 5 | 0.130 | 0.198 | 0.157 | 0.030 |
| | | Fractal dome | 5 | 0.177 | 0.235 | 0.211 | 0.028 |
| | | Control hexagon | 5 | 0.190 | 0.225 | 0.201 | 0.014 |
| | | Superdome | 8 | 0.115 | 0.241 | 0.164 | 0.040 |
| 6 months | Reef 13 | Control | 8 | 0 | 0.004 | 0.001 | 0.002 |
| | | Open hexagon | 5 | 0.004 | 0.011 | 0.007 | 0.003 |
| | | Closed hexagon | 5 | 0.005 | 0.014 | 0.008 | 0.004 |
| | | Fractal dome | 5 | 0.015 | 0.038 | 0.021 | 0.009 |
| | | Control hexagon | 5 | 0.018 | 0.024 | 0.020 | 0.002 |
| | | Sizedome | 8 | 0.010 | 0.023 | 0.017 | 0.004 |
| | | Superdome | 8 | 0.005 | 0.031 | 0.022 | 0.009 |
| | Moku o Loʻe | Control | 3 | 0.004 | 0.013 | 0.008 | 0.005 |
| | | Open hexagon | 5 | 0.015 | 0.043 | 0.032 | 0.011 |
| | | Closed hexagon | 5 | 0.058 | 0.086 | 0.072 | 0.011 |
| | | Fractal dome | 5 | 0.051 | 0.124 | 0.074 | 0.030 |
| | | Control hexagon | 5 | 0.029 | 0.054 | 0.044 | 0.010 |
| | | Superdome | 8 | 0.033 | 0.092 | 0.064 | 0.018 |
| 1 year | Reef 13 | Control | 8 | 0.000 | 0.002 | 0.001 | 0.001 |
| | | Open hexagon | 5 | 0.006 | 0.019 | 0.012 | 0.005 |
| | | Closed hexagon | 5 | 0.009 | 0.017 | 0.013 | 0.004 |
| | | Fractal dome | 5 | 0.014 | 0.032 | 0.018 | 0.008 |
| | | Control hexagon | 5 | 0.013 | 0.027 | 0.019 | 0.006 |
| | | Sizedome | 8 | 0.007 | 0.023 | 0.017 | 0.006 |
| | | Superdome | 8 | 0.011 | 0.023 | 0.020 | 0.004 |
| | Moku o Loʻe | Control | 3 | 0.002 | 0.004 | 0.004 | 0.001 |
| | | Open hexagon | 5 | 0.020 | 0.030 | 0.027 | 0.004 |
| | | Closed hexagon | 5 | 0.014 | 0.045 | 0.029 | 0.012 |
| | | Fractal dome | 5 | 0.020 | 0.049 | 0.032 | 0.012 |
| | | Control hexagon | 5 | 0.011 | 0.034 | 0.023 | 0.009 |
| | | Superdome | 8 | 0.026 | 0.050 | 0.037 | 0.008 |

**Table S5.**

Post hoc power analysis for all pairwise comparisons of dome types across timepoints based on total coral recruit numbers per module. Cohen's d (standardized effect size) and estimated statistical power are given for each comparison, calculated using the observed means, standard deviations, and sample sizes. Comparisons with power ≥ 0.8 indicate sufficient sensitivity to detect the observed effect sizes, comparisons with power < 0.8 may be underpowered.

| Timepoint | Group1 | Group2 | Cohen's d | Power |
|---|---|---|---|---|
| 1 week | Control dome | Open hexagon | 1.78 | 0.983 |
| | Control dome | Closed hexagon | 1.94 | 0.993 |
| | Control dome | Fractal dome | 1.14 | 0.744 |
| | Control dome | Hexagon dome | 1.12 | 0.729 |
| | Control dome | Sizedome | 5.84 | 1.000 |
| | Control dome | Superdome | 1.38 | 0.942 |
| | Open hexagon | Closed hexagon | 0.52 | 0.216 |
| | Open hexagon | Fractal dome | 0.86 | 0.489 |
| | Open hexagon | Hexagon dome | 0.90 | 0.522 |
| | Open hexagon | Sizedome | 5.42 | 1.000 |
| | Open hexagon | Superdome | 1.22 | 0.857 |
| | Closed hexagon | Fractal dome | 0.72 | 0.367 |
| | Closed hexagon | Hexagon dome | 0.79 | 0.423 |
| | Closed hexagon | Sizedome | 5.18 | 1.000 |
| | Closed hexagon | Superdome | 1.14 | 0.807 |
| | Fractal dome | Hexagon dome | 0.16 | 0.065 |
| | Fractal dome | Sizedome | 2.79 | 1.000 |
| | Fractal dome | Superdome | 0.62 | 0.335 |
| | Hexagon dome | Sizedome | 2.23 | 0.997 |
| | Hexagon dome | Superdome | 0.45 | 0.204 |
| | Sizedome | Superdome | 1.25 | 0.821 |
| 2 months | Control dome | Open hexagon | 1.30 | 0.846 |
| | Control dome | Closed hexagon | 1.42 | 0.900 |
| | Control dome | Fractal dome | 1.39 | 0.890 |
| | Control dome | Hexagon dome | 1.46 | 0.918 |
| | Control dome | Sizedome | 3.22 | 1.000 |
| | Control dome | Superdome | 1.69 | 0.991 |
| | Open hexagon | Closed hexagon | 0.67 | 0.323 |
| | Open hexagon | Fractal dome | 0.78 | 0.416 |
| | Open hexagon | Hexagon dome | 0.75 | 0.389 |
| | Open hexagon | Sizedome | 0.67 | 0.293 |
| | Open hexagon | Superdome | 1.05 | 0.741 |
| | Closed hexagon | Fractal dome | 0.18 | 0.068 |
| | Closed hexagon | Hexagon dome | 0.09 | 0.054 |
| | Closed hexagon | Sizedome | 1.09 | 0.635 |
| | Closed hexagon | Superdome | 0.40 | 0.166 |
| | Fractal dome | Hexagon dome | 0.09 | 0.055 |
| | Fractal dome | Sizedome | 1.13 | 0.661 |
| | Fractal dome | Superdome | 0.20 | 0.079 |
| | Hexagon dome | Sizedome | 1.16 | 0.684 |
| | Hexagon dome | Superdome | 0.31 | 0.120 |

| | | | | |
|---|---|---|---|---|
| | Sizedome | Superdome | 1.42 | 0.908 |
| 6 months | Control dome | Open hexagon | 1.57 | 0.948 |
| | Control dome | Closed hexagon | 1.55 | 0.943 |
| | Control dome | Fractal dome | 1.81 | 0.986 |
| | Control dome | Hexagon dome | 2.85 | 1.000 |
| | Control dome | Sizedome | 4.11 | 1.000 |
| | Control dome | Superdome | 2.30 | 1.000 |
| | Open hexagon | Closed hexagon | 0.92 | 0.535 |
| | Open hexagon | Fractal dome | 1.10 | 0.691 |
| | Open hexagon | Hexagon dome | 0.89 | 0.509 |
| | Open hexagon | Sizedome | 0.17 | 0.065 |
| | Open hexagon | Superdome | 1.56 | 0.973 |
| | Closed hexagon | Fractal dome | 0.08 | 0.054 |
| | Closed hexagon | Hexagon dome | 0.50 | 0.200 |
| | Closed hexagon | Sizedome | 0.90 | 0.475 |
| | Closed hexagon | Superdome | 0.42 | 0.180 |
| | Fractal dome | Hexagon dome | 0.65 | 0.305 |
| | Fractal dome | Sizedome | 1.10 | 0.639 |
| | Fractal dome | Superdome | 0.36 | 0.143 |
| | Hexagon dome | Sizedome | 1.01 | 0.570 |
| | Hexagon dome | Superdome | 1.12 | 0.791 |
| | Sizedome | Superdome | 1.59 | 0.957 |
| 1 year | Control dome | Open hexagon | 2.82 | 1.000 |
| | Control dome | Closed hexagon | 2.25 | 0.999 |
| | Control dome | Fractal dome | 2.73 | 1.000 |
| | Control dome | Hexagon dome | 3.67 | 1.000 |
| | Control dome | Sizedome | 3.89 | 1.000 |
| | Control dome | Superdome | 3.62 | 1.000 |
| | Open hexagon | Closed hexagon | 0.49 | 0.198 |
| | Open hexagon | Fractal dome | 0.68 | 0.329 |
| | Open hexagon | Hexagon dome | 0.27 | 0.092 |
| | Open hexagon | Sizedome | 0.33 | 0.109 |
| | Open hexagon | Superdome | 1.84 | 0.995 |
| | Closed hexagon | Fractal dome | 0.10 | 0.056 |
| | Closed hexagon | Hexagon dome | 0.33 | 0.113 |
| | Closed hexagon | Sizedome | 0.29 | 0.092 |
| | Closed hexagon | Superdome | 1.15 | 0.816 |
| | Fractal dome | Hexagon dome | 0.50 | 0.201 |
| | Fractal dome | Sizedome | 0.46 | 0.161 |
| | Fractal dome | Superdome | 1.12 | 0.794 |
| | Hexagon dome | Sizedome | 0.07 | 0.052 |
| | Hexagon dome | Superdome | 1.72 | 0.990 |
| | Sizedome | Superdome | 1.69 | 0.974 |

**Table S6.**

Differences in recruit densities (recruits per $cm^2$) between the different shapes (control, open hexagon, closed hexagon, fractal dome, hexagon dome, sizedome, and superdome) and natural substrate in the barrier reef in after two months. Results are derived from Wilcoxon tests followed by holm-adjustment for multiple testing. Number of replicates per test group (n1 and n2), test statistic ($chi^2$), and p-value are given with bold values indicating significance ($p < 0.05$).

| Contrast | | | n1 | n2 | Statistic | p |
|---|---|---|---|---|---|---|
| Control | - | Reef | 11 | 32 | 284 | 0.311 |
| Open hexagon | - | Reef | 10 | 32 | 542 | **<0.001** |
| Closed hexagon | - | Reef | 10 | 32 | 638 | **<0.001** |
| Fractal dome | - | Reef | 10 | 32 | 626 | **<0.001** |
| Control hexagon | - | Reef | 10 | 32 | 638 | **<0.001** |
| Sizedome | - | Reef | 8 | 32 | 512 | **<0.001** |
| Superdome | - | Reef | 16 | 32 | 1024 | **<0.001** |

**Table S7.**

Differences in recruit numbers (total numbers summarized from three replicate recesses of same depth on one dome) and recruit densities (number of recruits per $cm^2$) between the different recess depths (1, 2, 3, and 4 cm) after one week, two months, six months, and one year. Results are derived from Wilcoxon tests followed by holm-adjustment for multiple testing. Number of replicates per test group (n1 and n2), test statistic ($chi^2$), and p-value are given with bold values indicating significance ($p < 0.05$).

| Parameter | Timepoint | Contrast | n1 | n2 | Statistic | p |
|---|---|---|---|---|---|---|
| Recruit numbers (Total number of recruits) | 1 week | 1 - 2 | 8 | 8 | 2 | 0.0040 |
| | | 1 - 3 | 8 | 8 | 2 | 0.0040 |
| | | 1 - 4 | 8 | 8 | 4 | 0.0070 |
| | | 2 - 3 | 8 | 8 | 22.5 | 0.6880 |
| | | 2 - 4 | 8 | 8 | 31.5 | 1 |
| | | 3 - 4 | 8 | 8 | 46 | 0.4830 |
| | 2 months | 1 - 2 | 8 | 8 | 18 | 0.3250 |
| | | 1 - 3 | 8 | 8 | 15.5 | 0.3250 |
| | | 1 - 4 | 8 | 8 | 5 | **0.0300** |
| | | 2 - 3 | 8 | 8 | 27.5 | 0.6700 |
| | | 2 - 4 | 8 | 8 | 12 | 0.1940 |
| | | 3 - 4 | 8 | 8 | 15 | 0.3250 |
| | 6 months | 1 - 2 | 8 | 8 | 27 | 1 |
| | | 1 - 3 | 8 | 8 | 9.5 | 0.1170 |
| | | 1 - 4 | 8 | 8 | 11.5 | 0.1680 |
| | | 2 - 3 | 8 | 8 | 15.5 | 0.3560 |
| | | 2 - 4 | 8 | 8 | 16.5 | 0.3560 |
| | | 3 - 4 | 8 | 8 | 35.5 | 1 |
| | 1 year | 1 - 2 | 8 | 8 | 12.5 | 0.1640 |
| | | 1 - 3 | 8 | 8 | 7.5 | 0.0540 |
| | | 1 - 4 | 8 | 8 | 1 | **0.0070** |
| | | 2 - 3 | 8 | 8 | 20 | 0.4420 |
| | | 2 - 4 | 8 | 8 | 13 | 0.1640 |
| | | 3 - 4 | 8 | 8 | 27 | 0.6310 |
| Recruit density (Number of recruits per $cm^2$) | 1 week | 1 - 2 | 8 | 8 | 10 | 0.1040 |
| | | 1 - 3 | 8 | 8 | 15 | 0.2490 |
| | | 1 - 4 | 8 | 8 | 35 | 1 |
| | | 2 - 3 | 8 | 8 | 28 | 1 |
| | | 2 - 4 | 8 | 8 | 56 | 0.0620 |
| | | 3 - 4 | 8 | 8 | 53 | 0.1120 |
| | 2 months | 1 - 2 | 8 | 8 | 24 | 1 |
| | | 1 - 3 | 8 | 8 | 26 | 1 |
| | | 1 - 4 | 8 | 8 | 13 | 0.3010 |
| | | 2 - 3 | 8 | 8 | 34 | 1 |
| | | 2 - 4 | 8 | 8 | 22 | 1 |
| | | 3 - 4 | 8 | 8 | 18 | 0.7750 |
| | 6 months | 1 - 2 | 8 | 8 | 31 | 1 |
| | | 1 - 3 | 8 | 8 | 19 | 1 |
| | | 1 - 4 | 8 | 8 | 26 | 1 |
| | | 2 - 3 | 8 | 8 | 20 | 1 |
| | | 2 - 4 | 8 | 8 | 27 | 1 |
| | | 3 - 4 | 8 | 8 | 43 | 1 |
| | 1 year | 1 - 2 | 8 | 8 | 17 | 0.4960 |
| | | 1 - 3 | 8 | 8 | 11 | 0.1780 |
| | | 1 - 4 | 8 | 8 | 12 | 0.1940 |
| | | 2 - 3 | 8 | 8 | 26 | 1 |
| | | 2 - 4 | 8 | 8 | 29 | 1 |
| | | 3 - 4 | 8 | 8 | 32 | 1 |

**Table S8.**

Summary statistics of positions of recruits inside the recesses of superdome modules (specific width and depth in mm) and associated light levels (µmol $m^{-2}$ $s^{-1}$, PAR) at reef 13 after one week, two months, six months, and one year. The number of replicates (n), minimum (min), maximum (max), median, 1st (q1) and 3rd quartile (q3) are given with mean and standard deviation (sd).

| Parameter | Timepoint | n | min | max | median | q1 | q3 | mean | sd |
|---|---|---|---|---|---|---|---|---|---|
| Recess width (cm) | 1 week | 239 | 0.34 | 3.15 | 1.24 | 0.99 | 1.70 | 1.40 | 0.64 |
| | 2 months | 31 | 0.24 | 2.90 | 1.48 | 0.99 | 1.99 | 1.50 | 0.72 |
| | 6 months | 83 | 0.52 | 2.62 | 1.54 | 1.10 | 1.91 | 1.52 | 0.53 |
| | 1 year | 52 | 0.40 | 2.60 | 1.40 | 1.10 | 2.00 | 1.52 | 0.63 |
| Recess depth (cm) | 1 week | 239 | 0.38 | 4.06 | 1.95 | 1.29 | 2.50 | 1.91 | 0.82 |
| | 2 months | 31 | 0.41 | 3.43 | 1.72 | 0.70 | 2.50 | 1.70 | 0.95 |
| | 6 months | 83 | 0.51 | 3.08 | 1.70 | 1.11 | 2.21 | 1.69 | 0.70 |
| | 1 year | 52 | 0.38 | 3.50 | 1.73 | 1.13 | 2.28 | 1.74 | 0.79 |
| Light level | 1 week | 239 | 18.7 | 182.0 | 32.6 | 27.8 | 42.6 | 51.8 | 48.2 |
| (µmol $m^{-2}$ $s^{-1}$, PAR) | 1 months | 31 | 10.2 | 181.0 | 42.0 | 32.7 | 77.5 | 69.4 | 60.4 |
| | 6 months | 83 | 21.4 | 179.0 | 43.5 | 33.7 | 62.9 | 62.6 | 47.8 |
| | 1 year | 52 | 18.7 | 182.0 | 41.8 | 31.6 | 62.7 | 59.9 | 48.2 |

**Table S9.**

Differences in recruit numbers (total numbers summarized from three replicate recesses of same depth on one dome) and recruit densities (number of recruits per $cm^2$) between the different positions within the recesses (CE: cryptic edge, I: inside, EE: exposed edge, O: outside), split by recess depths (1, 2, 3, and 4 cm) after one week, two months, six months, and one year. Results are derived from Wilcoxon tests followed by holm-adjustment for multiple testing. Number of replicates per test group (n1 and n2), test statistic ($chi^2$), and p-value are given with bold values indicating significance ($p < 0.05$).

| Parameter | Timepoint | Contrast | | | n1 | n2 | statistic | p.adj |
|---|---|---|---|---|---|---|---|---|
| Recruit numbers (Total number of recruits) | 1 week | CE | - | EE | 8 | 8 | 64 | **0.0020** |
| | | CE | - | I | 8 | 8 | 64 | **0.0009** |
| | | CE | - | O | 8 | 8 | 64 | **0.0020** |
| | | EE | - | I | 8 | 8 | 4 | **0.0040** |
| | | EE | - | O | 8 | 8 | 28 | 0.3820 |
| | | I | - | O | 8 | 8 | 60 | **0.0050** |
| | | CE | - | EE | 8 | 8 | 64 | **0.0040** |
| | | CE | - | I | 8 | 8 | 63 | **0.0050** |
| | | CE | - | O | 8 | 8 | 64 | **0.0020** |
| | | EE | - | I | 8 | 8 | 11 | **0.0490** |
| | | EE | - | O | 8 | 8 | 44 | 0.0760 |
| | | I | - | O | 8 | 8 | 56 | **0.0140** |
| | | CE | - | EE | 8 | 8 | 64 | **0.0040** |
| | | CE | - | I | 8 | 8 | 54 | 0.0550 |
| | | CE | - | O | 8 | 8 | 64 | **0.0040** |
| | | EE | - | I | 8 | 8 | 0 | **0.0040** |
| | | EE | - | O | 8 | 8 | 33 | 0.9450 |
| | | I | - | O | 8 | 8 | 64 | **0.0040** |
| | | CE | - | EE | 8 | 8 | 64 | **0.0030** |
| | | CE | - | I | 8 | 8 | 58 | **0.0150** |
| | | CE | - | O | 8 | 8 | 64 | **0.0020** |
| | | EE | - | I | 8 | 8 | 0 | **0.0030** |
| | | EE | - | O | 8 | 8 | 40 | 0.1700 |
| | | I | - | O | 8 | 8 | 64 | **0.0020** |
| | 2 months | CE | - | EE | 8 | 8 | 3 | **0.0220** |
| | | CE | - | I | 8 | 8 | 1 | 0.2300 |
| | | CE | - | O | 8 | 8 | 3 | **0.0220** |
| | | EE | - | I | 8 | 8 | 2 | 0.0970 |
| | | EE | - | O | 8 | 8 | NA | NA |
| | | I | - | O | 8 | 8 | 2 | 0.0970 |
| | | CE | - | EE | 8 | 8 | 59 | **0.0200** |
| | | CE | - | I | 8 | 8 | 44 | 0.4820 |
| | | CE | - | O | 8 | 8 | 59 | **0.0200** |
| | | EE | - | I | 8 | 8 | 10 | 0.0500 |
| | | EE | - | O | 8 | 8 | 32 | 1 |
| | | I | - | O | 8 | 8 | 54 | 0.0500 |
| | | CE | - | EE | 8 | 8 | 63 | **0.0040** |
| | | CE | - | I | 8 | 8 | 50 | 0.1230 |
| | | CE | - | O | 8 | 8 | 64 | **0.0020** |
| | | EE | - | I | 8 | 8 | 10 | **0.0370** |
| | | EE | - | O | 8 | 8 | 36 | 0.3820 |
| | | I | - | O | 8 | 8 | 56 | **0.0170** |
| | | CE | - | EE | 8 | 8 | 62 | **0.0100** |
| | | CE | - | I | 8 | 8 | 58 | **0.0270** |
| | | CE | - | O | 8 | 8 | 64 | **0.0040** |
| | | EE | - | I | 8 | 8 | 26 | 0.5430 |

| | | | | | | | | |
|---|---|---|---|---|---|---|---|---|
| | | EE | - | O | 8 | 8 | 44 | 0.3180 |
| | | I | - | O | 8 | 8 | 48 | 0.1830 |
| | 6 months | CE | - | EE | 8 | 8 | 4 | **0.0019** |
| | | CE | - | I | 8 | 8 | 2 | 0.0504 |
| | | CE | - | O | 8 | 8 | 4 | **0.0019** |
| | | EE | - | I | 8 | 8 | 2 | 0.1520 |
| | | EE | - | O | 8 | 8 | NA | NA |
| | | I | - | O | 8 | 8 | 2 | 0.1520 |
| | | CE | - | EE | 8 | 8 | 56 | 0.0590 |
| | | CE | - | I | 8 | 8 | 56 | 0.0590 |
| | | CE | - | O | 8 | 8 | 60 | **0.0090** |
| | | EE | - | I | 8 | 8 | 32 | 1 |
| | | EE | - | O | 8 | 8 | 44 | 0.2240 |
| | | I | - | O | 8 | 8 | 44 | 0.2240 |
| | | CE | - | EE | 8 | 8 | 58 | **0.0230** |
| | | CE | - | I | 8 | 8 | 54 | 0.0510 |
| | | CE | - | O | 8 | 8 | 64 | **0.0020** |
| | | EE | - | I | 8 | 8 | 21 | 0.2240 |
| | | EE | - | O | 8 | 8 | 52 | **0.0350** |
| | | I | - | O | 8 | 8 | 56 | **0.0220** |
| | | CE | - | EE | 8 | 8 | 61 | **0.0100** |
| | | CE | - | I | 8 | 8 | 62 | **0.0080** |
| | | CE | - | O | 8 | 8 | 64 | **0.0020** |
| | | EE | - | I | 8 | 8 | 43 | 0.2500 |
| | | EE | - | O | 8 | 8 | 56 | **0.0110** |
| | | I | - | O | 8 | 8 | 44 | 0.1520 |
| | 1 year | CE | - | EE | 8 | 8 | 3 | **0.0069** |
| | | CE | - | I | 8 | 8 | 3 | **0.0099** |
| | | CE | - | O | 8 | 8 | 3 | **0.0069** |
| | | EE | - | I | 8 | 8 | 1 | 0.7631 |
| | | EE | - | O | 8 | 8 | NA | NA |
| | | I | - | O | 8 | 8 | 1 | 0.7631 |
| | | CE | - | EE | 8 | 8 | 58 | **0.0230** |
| | | CE | - | I | 8 | 8 | 61 | **0.0100** |
| | | CE | - | O | 8 | 8 | 64 | **0.0020** |
| | | EE | - | I | 8 | 8 | 38 | 0.5500 |
| | | EE | - | O | 8 | 8 | 48 | 0.0950 |
| | | I | - | O | 8 | 8 | 44 | 0.1490 |
| | | CE | - | EE | 8 | 8 | 58 | **0.0230** |
| | | CE | - | I | 8 | 8 | 57 | **0.0300** |
| | | CE | - | O | 8 | 8 | 60 | **0.0080** |
| | | EE | - | I | 8 | 8 | 22 | 0.2500 |
| | | EE | - | O | 8 | 8 | 44 | 0.1520 |
| | | I | - | O | 8 | 8 | 56 | **0.0190** |
| | | CE | - | EE | 8 | 8 | 61 | **0.0070** |
| | | CE | - | I | 8 | 8 | 64 | **0.0030** |
| | | CE | - | O | 8 | 8 | 64 | **0.0020** |
| | | EE | - | I | 8 | 8 | 55 | **0.0220** |
| | | EE | - | O | 8 | 8 | 60 | **0.0050** |
| | | I | - | O | 8 | 8 | 40 | 0.1700 |
| Recruit density (Number of recruits per cm$^2$) | 1 week | CE | - | EE | 8 | 8 | 64 | **0.0020** |
| | | CE | - | I | 8 | 8 | 64 | **0.0009** |
| | | CE | - | O | 8 | 8 | 64 | **0.0020** |
| | | EE | - | I | 8 | 8 | 4 | **0.0040** |
| | | EE | - | O | 8 | 8 | 28 | 0.3820 |
| | | I | - | O | 8 | 8 | 60 | **0.0050** |
| | | CE | - | EE | 8 | 8 | 64 | **0.0040** |
| | | CE | - | I | 8 | 8 | 63 | **0.0050** |

| | | | | | | | |
|---|---|---|---|---|---|---|---|
| | CE | - | O | 8 | 8 | 64 | **0.0020** |
| | EE | - | I | 8 | 8 | 11 | **0.0490** |
| | EE | - | O | 8 | 8 | 44 | 0.0760 |
| | I | - | O | 8 | 8 | 56 | **0.0140** |
| | CE | - | EE | 8 | 8 | 64 | **0.0040** |
| | CE | - | I | 8 | 8 | 30 | 1 |
| | CE | - | O | 8 | 8 | 64 | **0.0040** |
| | EE | - | I | 8 | 8 | 0 | **0.0040** |
| | EE | - | O | 8 | 8 | 32 | 1 |
| | I | - | O | 8 | 8 | 64 | **0.0040** |
| | CE | - | EE | 8 | 8 | 64 | **0.0030** |
| | CE | - | I | 8 | 8 | 18 | 0.3120 |
| | CE | - | O | 8 | 8 | 64 | **0.0020** |
| | EE | - | I | 8 | 8 | 0 | **0.0030** |
| | EE | - | O | 8 | 8 | 40 | 0.3120 |
| | I | - | O | 8 | 8 | 64 | **0.0020** |
| 2 months | CE | - | EE | 8 | 8 | 3 | **0.0220** |
| | CE | - | I | 8 | 8 | 2 | 0.0970 |
| | CE | - | O | 8 | 8 | 3 | **0.0220** |
| | EE | - | I | 8 | 8 | 2 | 0.0970 |
| | EE | - | O | 8 | 8 | NA | NA |
| | I | - | O | 8 | 8 | 2 | 0.0970 |
| | CE | - | EE | 8 | 8 | 58 | **0.0250** |
| | CE | - | I | 8 | 8 | 40 | 0.8560 |
| | CE | - | O | 8 | 8 | 60 | **0.0140** |
| | EE | - | I | 8 | 8 | 9 | **0.0360** |
| | EE | - | O | 8 | 8 | 33 | 1 |
| | I | - | O | 8 | 8 | 55 | **0.0360** |
| | CE | - | EE | 8 | 8 | 62 | **0.0060** |
| | CE | - | I | 8 | 8 | 40 | 0.7640 |
| | CE | - | O | 8 | 8 | 64 | **0.0020** |
| | EE | - | I | 8 | 8 | 9 | **0.0270** |
| | EE | - | O | 8 | 8 | 36 | 0.7640 |
| | I | - | O | 8 | 8 | 56 | **0.0170** |
| | CE | - | EE | 8 | 8 | 61 | **0.0120** |
| | CE | - | I | 8 | 8 | 41 | 0.5400 |
| | CE | - | O | 8 | 8 | 62 | **0.0070** |
| | EE | - | I | 8 | 8 | 20 | 0.5400 |
| | EE | - | O | 8 | 8 | 43 | 0.5400 |
| | I | - | O | 8 | 8 | 49 | 0.2130 |
| 6 months | CE | - | EE | 8 | 8 | 4 | **0.0019** |
| | CE | - | I | 8 | 8 | 3 | **0.0067** |
| | CE | - | O | 8 | 8 | 4 | **0.0019** |
| | EE | - | I | 8 | 8 | 2 | 0.1520 |
| | EE | - | O | 8 | 8 | NA | NA |
| | I | - | O | 8 | 8 | 2 | 0.1520 |
| | CE | - | EE | 8 | 8 | 53 | 0.1520 |
| | CE | - | I | 8 | 8 | 53 | 0.1520 |
| | CE | - | O | 8 | 8 | 60 | **0.0090** |
| | EE | - | I | 8 | 8 | 28 | 0.6270 |
| | EE | - | O | 8 | 8 | 44 | 0.2240 |
| | I | - | O | 8 | 8 | 44 | 0.2240 |
| | CE | - | EE | 8 | 8 | 54 | 0.0670 |
| | CE | - | I | 8 | 8 | 45 | 0.1860 |
| | CE | - | O | 8 | 8 | 64 | **0.0020** |
| | EE | - | I | 8 | 8 | 15 | 0.1500 |
| | EE | - | O | 8 | 8 | 52 | **0.0470** |
| | I | - | O | 8 | 8 | 56 | **0.0220** |

| | | | | | | | |
|---|---|---|---|---|---|---|---|
| | CE | - | EE | 8 | 8 | 58 | **0.0260** |
| | CE | - | I | 8 | 8 | 57 | **0.0270** |
| | CE | - | O | 8 | 8 | 64 | **0.0020** |
| | EE | - | I | 8 | 8 | 35 | 0.7800 |
| | EE | - | O | 8 | 8 | 56 | **0.0190** |
| | I | - | O | 8 | 8 | 44 | 0.1520 |
| 1 year | CE | - | EE | 8 | 8 | 3 | **0.0069** |
| | CE | - | I | 8 | 8 | 3 | **0.0070** |
| | CE | - | O | 8 | 8 | 3 | **0.0069** |
| | EE | - | I | 8 | 8 | 1 | 0.7631 |
| | EE | - | O | 8 | 8 | NA | NA |
| | I | - | O | 8 | 8 | 1 | 0.7631 |
| | CE | - | EE | 8 | 8 | 54 | 0.0880 |
| | CE | - | I | 8 | 8 | 58 | **0.0310** |
| | CE | - | O | 8 | 8 | 64 | **0.0020** |
| | EE | - | I | 8 | 8 | 33 | 0.9540 |
| | EE | - | O | 8 | 8 | 48 | 0.0950 |
| | I | - | O | 8 | 8 | 44 | 0.1490 |
| | CE | - | EE | 8 | 8 | 58 | **0.0270** |
| | CE | - | I | 8 | 8 | 55 | **0.0470** |
| | CE | - | O | 8 | 8 | 60 | **0.0080** |
| | EE | - | I | 8 | 8 | 16 | 0.1520 |
| | EE | - | O | 8 | 8 | 44 | 0.1520 |
| | I | - | O | 8 | 8 | 56 | **0.0190** |
| | CE | - | EE | 8 | 8 | 59 | **0.0150** |
| | CE | - | I | 8 | 8 | 62 | **0.0070** |
| | CE | - | O | 8 | 8 | 64 | **0.0020** |
| | EE | - | I | 8 | 8 | 47 | 0.2160 |
| | EE | - | O | 8 | 8 | 60 | **0.0070** |
| | I | - | O | 8 | 8 | 40 | 0.2160 |

**Table S10.**

Summary statistics of coral size and biomass parameters (diameter per recruit, planar surface area per recruit, volume per recruit, total planar surface area per dome, and total volume per dome) on control domes and superdomes at reef 13 and Moku o Lo‘e one year. The number of replicates (n), minimum (min) and maximum (max) values are given with mean and standard deviation (sd).

| Parameter | Location | Dome type | n | min | max | mean | sd |
|---|---|---|---|---|---|---|---|
| Diameter (mm) | Reef 13 | Control | 14 | 1 | 4 | 2.29 | 0.91 |
| | | Superdome | 104 | 1 | 11 | 3.40 | 1.91 |
| | Moku o Lo‘e | Control | 10 | 2 | 6 | 3.70 | 1.49 |
| | | Superdome | 272 | 1 | 12 | 2.89 | 1.63 |
| Planar surface area ($mm^2$) | Reef 13 | Control | 7 | 0.79 | 9.43 | 4.38 | 2.97 |
| | | Superdome | 52 | 0.79 | 55.00 | 11.40 | 12.40 |
| | Moku o Lo‘e | Control | 5 | 3.14 | 28.30 | 12.30 | 10.10 |
| | | Superdome | 136 | 0.79 | 104.00 | 8.38 | 11.50 |
| Volume ($mm^3$) | Reef 13 | Control | 7 | 0.79 | 18.80 | 7.40 | 6.90 |
| | | Superdome | 52 | 0.79 | 220.00 | 29.50 | 47.80 |
| | Moku o Lo‘e | Control | 5 | 6.28 | 113.00 | 43.50 | 44.60 |
| | | Superdome | 136 | 0.79 | 933.00 | 23.70 | 83.20 |
| Total planar surface area ($mm^2$) | Reef 13 | Control | 3 | 3.14 | 19.60 | 10.20 | 8.49 |
| | | Superdome | 3 | 156.00 | 264.00 | 197.00 | 58.20 |
| | Moku o Lo‘e | Control | 3 | 14.10 | 28.30 | 20.40 | 7.20 |
| | | Superdome | 5 | 144.00 | 281.00 | 228.00 | 54.20 |
| Total volume ($mm^3$) | Reef 13 | Control | 3 | 3.14 | 36.10 | 17.30 | 17.00 |
| | | Superdome | 3 | 448.00 | 579.00 | 512.00 | 65.30 |
| | Moku o Lo‘e | Control | 3 | 35.30 | 113.00 | 72.50 | 39.00 |
| | | Superdome | 5 | 250.00 | 1275.00 | 645.00 | 385.00 |

**Table S11.**

Differences in coral size and biomass parameters (diameter per recruit, planar surface area per recruit, volume per recruit, total planar surface area per dome, and total volume per dome) on control domes and superdomes at reef 13 and Moku o Lo‘e one year. Results are derived from Wilcoxon tests. Number of replicates per test group (n1 and n2), test statistic (chi2), and p-value are given with bold values indicating significance ($p < 0.05$).

| Parameter | Location | Contrast | | | n1 | n2 | statistic | p.adj |
|---|---|---|---|---|---|---|---|---|
| Diameter (mm) | Reef 13 | Control | - | Superdome | 14 | 104 | 473 | **0.0303** |
| | Moku o Lo‘e | Control | - | Superdome | 10 | 272 | 1860 | **0.0389** |
| Planar surface area ($mm^2$) | Reef 13 | Control | - | Superdome | 7 | 52 | 111 | 0.0959 |
| | Moku o Lo‘e | Control | - | Superdome | 5 | 136 | 467 | 0.1470 |
| Volume ($mm^3$) | Reef 13 | Control | - | Superdome | 7 | 52 | 124 | 0.1760 |
| | Moku o Lo‘e | Control | - | Superdome | 5 | 136 | 506 | 0.0639 |
| Total planar surface area ($mm^2$) | Reef 13 | Control | - | Superdome | 3 | 3 | 0 | 0.1000 |
| | Moku o Lo‘e | Control | - | Superdome | 3 | 5 | 0 | **0.0357** |
| Total volume ($mm^3$) | Reef 13 | Control | - | Superdome | 3 | 3 | 0 | 0.1000 |
| | Moku o Lo‘e | Control | - | Superdome | 3 | 5 | 0 | **0.0357** |

**Table S12.**

Differences in recruit densities between the different flow treatments (no, low, and high) on superdomes and control domes after one week, two months, six months, and one year. Results are derived from Wilcoxon test, followed by holm-adjustment for multiple testing (one week). Number of replicates per test group (n1 and n2), test statistic ($chi^2$ and z-statistic, respectively), and p-value are given with bold values indicating significance ($p < 0.05$).

| Timepoint | Dome type | Contrast | n1 | n2 | statistic | p |
|---|---|---|---|---|---|---|
| 1 week | Superdome | no - low | 7 | 8 | 8 | **0.0410** |
| | | no - high | 7 | 8 | 4 | **0.0110** |
| | | low - high | 8 | 8 | 17 | 0.1300 |
| | Control | no - low | 6 | 6 | 16 | 1.0000 |
| | | no - high | 6 | 6 | 15 | 1.0000 |
| | | low - high | 6 | 6 | 27 | 0.5400 |
| 2 months | Superdome | low - high | 11 | 12 | 83 | 0.3164 |
| | Control | low - high | 9 | 9 | 17 | **0.0379** |
| 6 months | Superdome | low - high | 11 | 12 | 74 | 0.6505 |
| | Control | low - high | 9 | 9 | 24 | 0.1615 |
